# The αβTCR repertoire at scale in the immgenT dataset

**DOI:** 10.64898/2026.01.30.702900

**Authors:** Myriam Croze, Liang Yang, Serge Candéias, Ian Magill, Odhran Casey, Val Piekarsa, Brinda Vijaykumar, Véronique Giudicelli, Sofia Kossida, David Zemmour, Christophe Benoist, the immgenT Project

## Abstract

The immense T cell receptor (TCR) repertoire is shaped by VDJ combinatorial diversity, imprecise rearrangements, and clonal selection. The immgenT Project generated scRNA and TCRseq to map paired αβTCR repertoires across 734 mouse samples from diverse tissues and challenge conditions. Compositional analysis uncovered some extreme junctional architectures. Beyond probabilistic V and J pairing, over-represented joins suggested non-randomness in fine joining, broadening the precedent of quasi-invariant iNKT and MAIT TCRs. We charted public clonotypes linked to self or environmental antigens in the main lineages. Tissue analyses revealed compartmentalized tissue-specific expansions. Unproductive and productive rearrangements of a V gene appeared to interfere specifically with each other, at chromatin or RNA levels. Unexpectedly, allelic exclusion at TCRβ proved less stringent than thought, and we identified rearrangements of TCRα in immature pre-T stages. This organism-wide look into the TCR repertoire offers novel insights on the evolutionary and immunological pressures on TCR repertoire selection.

## INTRODUCTION

The epicenter of T cell biology is the T-cell receptor (TCR), which drives or conditions virtually every moment of a T cell’s life, by binding short peptides presented by molecules encoded in the Major Histocompatibility Complex (pMHC). The TCRαβ heterodimer is encoded by randomly rearranged Variable, Diversity and Joining elements (*TRBV*, *TRBD* and *TRBJ* for TCRβ, and *TRAV* and *TRAJ* segments for TCRα). Variability arises from this combinatorial composition from large arrays of elements, amplified by random base insertions (N nucleotides) and deletions during the joining process^1,2^, yielding a potentially enormous number of clonotypes. Theoretical estimates ranging from 10^13^ to 10^20^ have been proposed^3–7^, dwarfing the number of T cells in a mouse (∼10^8^) or human (10^12^). Such numbers imply that every individual would sample only a small and different fraction of the potential “species repertoire”. These early calculations were based on the assumption that the combinatorial composition of genomic elements into productive TCR assemblies is random, However, probabilities of rearrangement vary widely for different V or J elements, as do the extent of nucleotide removal or addition. The repertoire is further constrained and made uneven by selection steps in the thymus (preTα selection, TCRαβ pairing, compatibility with MHC molecules) and clonal survival and expansion in the periphery.

How this highly diverse TCR repertoire is generated, is rendered [mostly] tolerant of self, supports T cell homeostasis, and is deployed with extensive amplification during responses to foreign antigens, is one of the most fascinating questions in adaptive immunology. The breadth of the TCR repertoire makes its study daunting, however. The first single-cell studies from the 1990s could only tackle tens to hundreds of cells^8–10^, an infinitesimal fraction of the repertoire. High-throughput sequencing allowed millions of cells to be tackled, but could only yield data for only one TCR chain at a time, giving a truncated view of the true repertoire^11^. Single-cell RNA/TCRseq technologies then allowed the determination of paired TCRαβ for thousands to hundreds of thousands of cells, bringing vaster perspectives on the repertoire^7,12,13^.

The immgenT program^14^, because of the vast array of lineages, tissues and challenged conditions in the 734 samples it profiled, provided the opportunity to analyze the T cell repertoire not solely from blood or secondary lymphoid organs but from a much larger organismal perspective, probing how thymic and peripheral clonal selection operate, and in a context where the clonotype analysis was enriched by information on each cell’s surface phenotype and transcriptome. This depth also brought a number of surprising observations, including unexpected structures created by rearrangements, effects of unproductive joins, recurring rearrangements and a perspective on how germline selection may balance the affinity between TCR and MHC molecules.

Companion web resources support public browsing and interrogation of the TCR data (https://www.immgen.org/ImmGenT/).

## RESULTS

The immgenT study analyzed altogether 680,000 T cells, mostly from mice on the C57Bl/6J background, encompassing 46 different anatomical sites and a broad array of immunological conditions (baseline, physiological states, infections, tumors and autoimmunity; **Fig. 1A**). Accompanying manuscripts detail the transcriptional characteristics across lineages and clusters. Single-cell RNAseq (scRNAseq) on the 10X Genomics 5’ v3 platform was used, deriving TCR-VDJ sequencing libraries that yielded paired αβTCR information on 418,463 cells (62% efficiency overall) for which single-cell transcriptomes were also available. TCR composition was determined from assemblies (CellRanger contigs) mapped at allele-level with IMGT/HighV-QUEST (V-QUEST), using a C57Bl/6-specific reference (this reference was corrected based on early immgenT data). This highly varied dataset yielded very rich information on TCR diversity across T cell lineages, as illustrated by the TCR-dist plot of **Fig. 1B**: the tight clustering of iNKT TCRs among Zbtb16+ nonconventional T cells (Tz), the TCRs of many CD8aa T cells that occupy a unique swath, the TCRs of CD4+, CD8+ and Treg cells distributed across several regions.

**Fig. 1.**
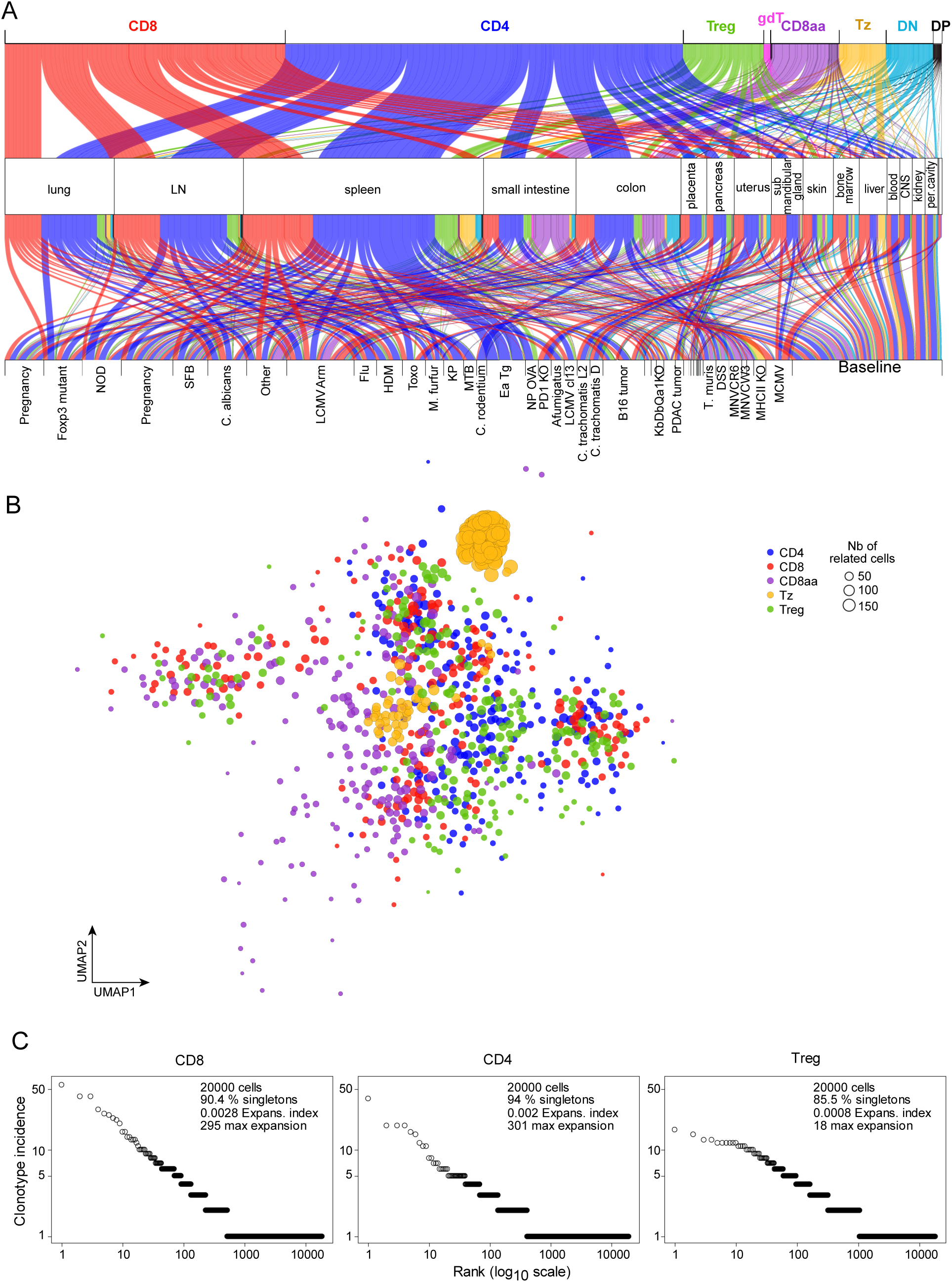
Overview of the immgenT TCR data. **A.** Single-cell TCRseq data from the immgenT program was generated in 66 experiments, originating from 46 organismal locations and >40 different conditions of immunological challenges. T cells encompassed in this analysis were grouped into main lineages (top row) as detailed in (immgenT-Cosmo) (top row; “Tz” refers to non-conventional T cells expressing the ZBTB16 (aka PLZF) transcription factor). The second and third rows display organs and conditions (baseline refers to unchallenged mice). **B.** Schematic overview of the TCRαβ repertoire, as approximated by the CoNGA tool, which generates a graph of clusters of related TCR clonotypes based on sequence distance between VDJ and junction sequences, and supported by similarities in single-cell gene expression. This network was computed with CoNGA from a 30,000 cell down-sample of the immgenT data, and each node is color-coded according to the annotation of its member cells. **C.** TCRαβ clones (full nucleotide-level identity, both chains) present in each of the three main lineages, ranked according to their total incidence across the non-transgenic immgenT dataset. Data for CD4+ and CD8+ T cells are downsampled to match Treg numbers. The “expansion index” is the incidence of most frequent clone divided by the size of the dataset for the lineage, “max expansion” is the number of cells expressing from the most frequent clone in each lineage.

The immgenT dataset comprises 91 different preparations and runs (“IGTx” identifiers), each including 4-10 samples identified by hashtagging, and corresponding to one or a few mice. In the terminology used here: a “clone” represents a group of cells from the same mouse that exactly share TCRαβ sequence (both α and β chains, junction identity at nucleotide level); because cells with complete identity are very rare between individuals, such identity within an individual is considered to reflect expansion from the same clonal precursor. A “clonotype” is defined as identical paired TCRαβ amino-acid sequences (identical V, J and junctional regions), “alpha-clonotype” and “beta-clonotype” representing amino-acid identity on only one chain. When dealing with rearrangement mechanics, “aN-clonotype” and “bN-clonotype” are used to denote identity at the nucleotide level. Finally, “meta-clonotypes” are closely related TCRs that determine similar T cell reactivity and fate (e.g. iNKT receptors). Analyses in the following sections use either the whole dataset (when clonal expansion is relevant) or often a subset thereof in which repeated clones within a sample were collapsed to one representative, to avoid skewing distributions. In the course of this work, we noted some artefacts with TCR data and its processing (detailed in **Supplementary Note**) which may be generally useful in interpreting TCRseq data. The integration of gene expression data into clusters that subsume cell-states across tissues and challenges is described fully in accompanying manuscripts (**cosmoT, CD8, Treg, CD4 papers**)

### Diversity across lineages

The random and imprecise rearrangements of TCRα- and TCRβ-encoding genes, and their pairing to generate the complete TCR expressed at the cell surface, entail a huge combinatorial diversity. Standard tools from population biology have been applied to estimate repertoire size from small datasets, but their limitations have been pointed out: the distribution of rare clones cannot be adequately deduced from those of larger ones, which reflect different homeostatic drives (maintenance versus clonal expansion)^6,15,16^. Thus, we refrained from attempting to estimate “repertoire size”, but instead compared the degree of clonal expansion in different lineages in the non-transgenic moiety of the immgenT dataset (**Fig. 1C**). For conventional CD4+ and CD8+ T cells, the incidence of expanded clonotypes followed a power law distribution: few highly expanded clones, long tail of singletons (90+%), as already noted in single-chain or bulk repertoires ^6,17^. These strong expansions result from antigen-driven responses, and are found in antigen-experienced cells (see accompanying **immgen-T Cosmology**). Interestingly, this distribution also applied to TCRs in Treg cells, with a. shallower curve: expansions of smaller sizes, none reaching the 200+ cell expansions of CD4+ or CD8+ cells, and a lower proportion of singletons. Although clonal expansions have been described in Treg cells ^18–21^, it wasn’t obvious that they would be such a common feature of the Treg pool, suggesting a narrower selection and/or more clonal expansion for Treg than for conventional T cells.

In addition, one of the immgenT datasets directly compared repertoires from adult and aged mice. As expected^22,23^, we observed a marked increase in the representation of expanded clonotypes in aged mice (**Fig. S1**)

### A compendium of public clonotypes

“Public TCRs” are clonotypes that appear recurrently in different individuals of a species. Their existence has been recognized for some time^24,25^, particularly in responses to viral infection^25–27^. They are generally attributed to the combination of high-probability rearrangements and antigen-driven selection^26,28^. Schematically, identities at the protein level denote functional selection, irrespective of the underlying sequence composition, while nucleotide-level identities denote high-probability rearrangements. The immgenT dataset allowed us to assess the distribution of public TCRs involving both chains of αβTCRs under a wide set of conditions, and to provide a reference set that can be of use as reference (https://rstats.immgen.org/tcrbrowser).

Among the 330,000 non-transgenic cells with full TCR data, we observed 682 αβTCR clonotypes that were shared between 2 to 8 independent datasets (**Fig. 2A, Table S1A**). These accounted for 3,254 cells altogether, ∼1% of the cells profiled. All lineages were represented (CD4+, CD8+, Treg, Tz). Among the latter, the quasi-invariant TCRs of iNKT cells (TRAV11.TRAJ18.CVVGDRGSALGRLHF) or MAIT cells (TRAV1.TRAJ33.CAVRDSNYQLIW) accounted for 221 of the public TCRs, with very little clonal expansion **(Table S1B**).

**Fig. 2.**
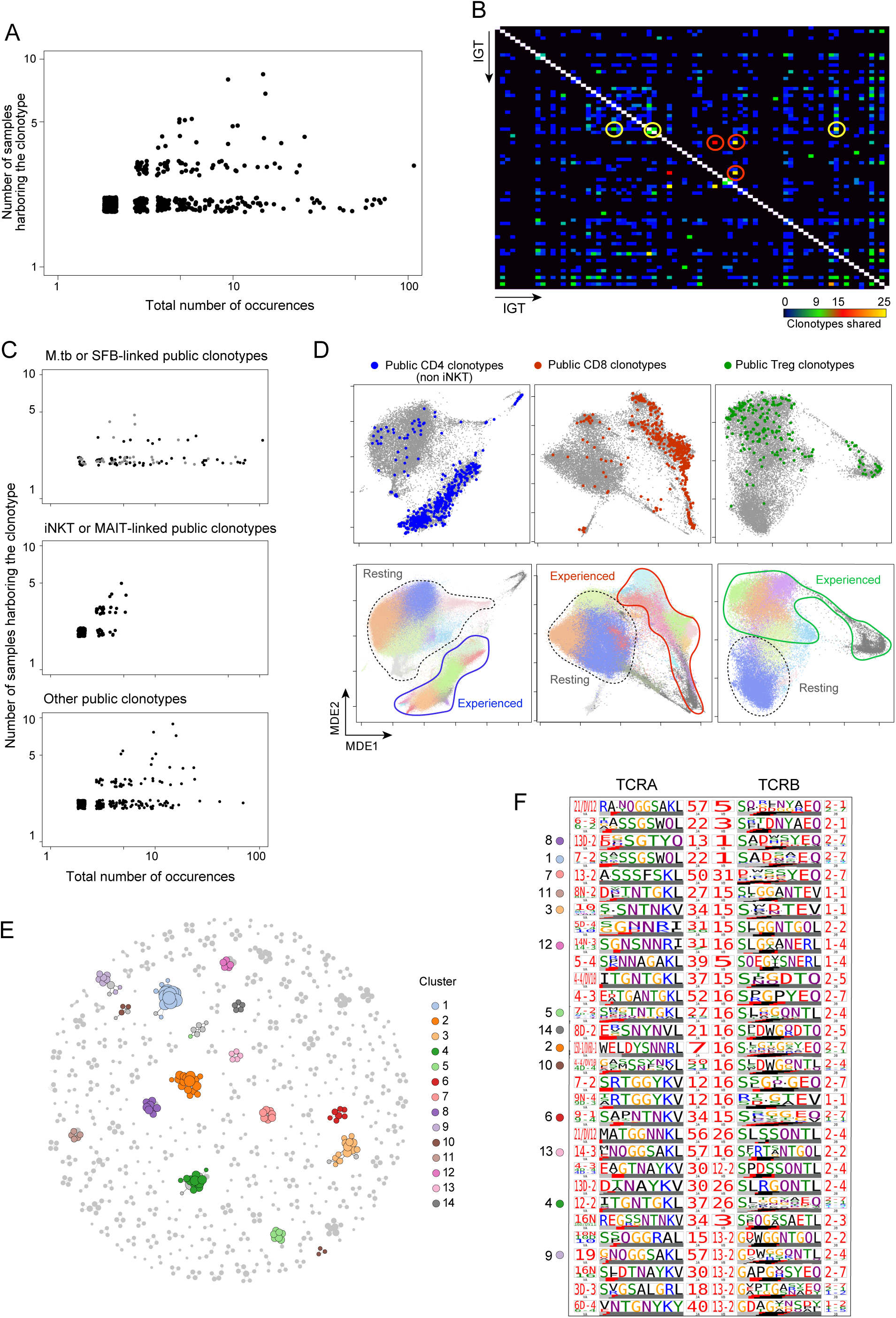
Public clonotypes in the immgenT data. Public clonotypes were first identified as TCRs that strictly share the same V, J and junction amino-acid sequences, across different donor mice (to ensure independence, across different experiments). **A.** Total number of occurrences across the immgenT dataset versus the number of independent experiments in which they appear. Each dot is an independent clonotype. **B.** Distribution of the sharing of public clonotypes between pairs of independent datasets (“IGT”). The more abundant sharing of public TCRs between IGTs from mice infected with SFB or *M. tuberculosis* (*M. tb*) are highlighted with yellow or red circles, respectively. **C.** As in **A,** but public clonotypes were separated between those linked to SFB and *M. tb* infection (top), quasi-invariant TCRs from iNKT and MAIT cells (middle) and “other” TCRs more generally fund in different lineages and not apparently related to any infection (see **Table S1**). **D.** T cells that express the “other” public clonotypes from **C** are displayed in color on the integrative immgenT MDE plots for CD4+, CD8+ and Treg lineages (top panels). For reference, immgenT clusters, and their partition into “resting” and “activated” clusters is shown below. **E.** The CoNGA algorithm was applied to 19,965 Treg cells assembled across 74 IGTs to identify public clonotypes defined by close similarity between TCR clonotypes, rather than by strict sequence identity. The TCR-distance plot highlights the major CoNGA clusters brought forth by the analysis (listed in **Table S2**). **F.** Sequence logos corresponding to the CoNGA clusters in **E**.

For 115 of these public TCRs, mice from different IGTs shared exactly the same TRA and TRB junctional nucleotide sequences. These had lower numbers of N additions (mean N additions in public joins =0.88 vs 2.49 in the overall TRA data, p<10^-18^), consistent with higher probability of generation, which was confirmed with the OLGA algorithm^29^ that predicts the likelihood of a particular VDJ join based on its composition (median Pgen for TCRα= 3.4x10^-6^ for these public TCRs vs 4.2x10^-7^ across the whole dataset, for TCRβ= 6.6x10^-8^ vs 4.2x10^-9^, Wilcoxon pval<10^-32^).

Sharing of public TCRs was widely distributed across IGT datasets (**Fig. 2B**), but some IGT pairs stood out with higher numbers of shared clonotypes (circled). These corresponded to microbe-related responses: one group was present in mice carrying the intestinal Segmented Filamented Bacterium (SFB), a pathobiont which elicits a strong Th17-biased response^30^ (**Table S1C**). SFB-positive mice from IGT37 and IGT90 shared 37 clonotypes, some of which correspond to TCRs discovered earlier in the Littman group (see accompanying **immgenT-Pregnancy**). Another 64 public TCRs were found in lung CD8+ and CD4+ T cells of *Mycobacterium tuberculosis (M. tb)*-infected mice (IGT47/61/66) (**Table S1D**). We are not suggesting that public TCRs are preferentially elicited by these two infectious agents relative to others, it is rather that their chance of discovery is higher because the immgenT data includes more samples related to these two microbes, increasing the odds of discovery.

Other than these iNKT/MAIT- or infection-related public TCRs, another set of 355 public clonotypes were present in 1,486 cells across many IGTs and in a variety of tissues, and were lineage-specific (**Table S1E**). These “other” public clonotypes showed more clonal expansion than those of iNKT/MAIT cells (**Fig. 2C**). When projected onto the integrated lineage maps derived from the gene expression data (**CosmoT manuscript**), public clonotypes were mostly found in antigen-experienced cell states (**Fig. 2D**), suggesting reactivity to common self or environmental antigens. Overall, public TCRs are thus quite prevalent, with ∼0.1% of clonotypes that occur repeatedly in different mice at baseline, and from which immune responses expand an array of microbe-specific recurring TCRs. Of course, we analyzed here genetically homogeneous B6 mice, and variable MHCs in outbred species would reduce this frequency.

These public TCRs were identified strictly, from complete sequence identity on both chains, but TCRs with small differences in CDR3 sequence can have the same reactivity^31^. Thus, the space of public TCRs should also include fuzzier meta-clonotypes. To identify these, we used the TCRdist algorithm in the CoNGA suite^32^, which scores biochemically-informed similarities between TCRs. Several “Conga clusters” of TCRs with closely related sequences were discovered in this manner, as exemplified for Treg cells in **Fig. 2E-F**, with 14 clusters that each included 5 to 29 clonotypes at FDR<0.01, across 19,895 Treg cells (note that CoNGA clusters do not overlap with strict public TCRs, as the algorithm collapses clones). All of these clusters were broadly represented across datasets, but they exhibited distinct tissue preferences (clusters 1, 2 and 8 mostly in gut, 5 in lung, 4 and 7 more widespread, **Table S2**). Thus, between public clonotypes and CoNGA clusters, the TCR repertoire includes many shared specificities that are found recurrently across individuals. It should also be stated that, because our sampling was shallow relative to the overall T cell pool of a mouse, we only detected the tip of the iceberg of public TCRs, those that are revealed by activation and amplification by common antigens, whether infectious (*M. tb* and SFB) or self/environmental (“other” group).

### Distribution of the TCR repertoire in single individuals

How T cells circulate between SLOs and parenchymal organs is of keen current interest. Most studies of the TCR repertoire encompass T cells in lymphoid organs (blood for human studies), or focus on one specific organ. Here, in “one-mouse” experiments, we assessed the distribution of TCR clonotypes across different locations of a single mouse, asking how widely amplified clonotypes disseminate within or between organs (**Fig. 3A**; another experiment was performed on a second mouse, which reproduced all the points made below, **Fig. S2A, B**). Total CD3+ T cells were profiled, yielding 16,242 cells with productive TCRα and TCRβ pairs. As illustrated in **Fig. 3B**, diversity was very different for SLOs and tissues (Gini indices around 0.001 for the former, 0.02 to 0.16 for the latter), with 25-30% of repeated clonotypes (among a uniform sampling of 100 TCRs) in the skin or kidney. These distributions yield a predicted repertoire size of 1-2x10^6^ for SLOs, much lower for tissues (**Fig. 3C**; Chao1 estimates are not valid when such clonal expansions are present). Grouping all the tissues, we detected 266 amplified clonotypes (defined by nucleotide-level TCRα and TCRβ identity) present in 944 cells. These were mostly private specificities, not broadly represented across the entire immgenT dataset. The distribution of these amplified clonotypes across the different tissues (**Fig. 3D, Table S3A-B**), revealed several interesting points:

i. Unexpectedly, most expanded clonotypes were found among kidney-associated T cells. B6 mice are not especially prone to kidney disease, compared to nephropathy-susceptible inbred strains. These expanded clonotypes involved a diversity of V regions (**Table S3A-B**), with no obvious convergence of CDR3 length or composition. They were found in CD4+ or CD8+ T cells, but comparatively fewer Tregs. A sizeable fraction of these kidney-preferential clonotypes (37.0%) was also found in the lung and liver of this mouse, and some in the blood, suggesting that this population can circulate widely (see also **Fig. S2B**). Cells carrying these expanded clonotypes distributed across several activated subclusters (**Fig. 3E**). Among CD4+ cells, clusters CD4-Q and CD4-S are mostly Th1-like, cluster CD4-I includes CD73+Izumo1r+ chronically stimulated cells – **see immgenT-CD4 manuscript**). Of the expanded CD8+ T cells, those in the kidney were dominated by those linked to chronic exposure and exhaustion (CD8.R/Q) and proliferating cells, while those in blood and other organs predominantly belonged to the CD8-H, more reminiscent of terminally-differentiated effector/memory state (**Fig. S2C, D**). This difference suggested that the kidney is likely where these T cells amplify in response to driving antigens (self or foreign?), with changes in cell states upon migration.
ii. Amplified clonotypes found in the intestines were completely distinct from the kidney group. In the lamina propria, amplified clonotypes were predominantly represented in Treg cells and CD4+ T cells (72.3 and 31.9%, respectively, **Fig. 3D**). There was very little overlap between clonotypes found in the intra-epithelial (IEL) and lamina propria compartments, supporting the notion that these are different compartments with limited exchange^33^. Some clonal sharing was found between the colon and small intestine (SI) lamina propria (Szymkiewicz–Simpson overlap coefficient=0.31, on a scale from 0 to 1), consistent with the noted exchange between different sections of the gut^34^, via draining mesenteric lymph node (mLN) or systemic circulation. On the other hand, sharing between colon and SI appeared much lower for IELs (**Fig. 3D**). This observation was confirmed in an independent IEL dataset (two B6 mice, profiling IELs from different regions of the gut, **Fig. 3F**). Expanded clonotypes were observed for the three types of αβTCR+ IELs (CD4+, CD8+, CD8aa+), and were shared between duodenum, jejunum and ileum, suggesting that IELs can traffic within the SI. Again, very few of these were found in colon IELs (**Fig. 3F**).
iii. Skin T cells also behaved as a distinct pool, with little/no overlap with other organs, but some exchange of clones between back and ear skin, in keeping with the notion that skin T cells are a specialized population that circulates little^35^.
iv. Pertinent to explorations of human diseases, for which blood is often the only accessible tissue, blood T cells included a variable representation of tissue-amplified clonotypes (**Fig. 3D, S2B**). The kidney/liver/lung clonotype group was represented in the blood, as were a few of the skin clonotypes, but virtually none of the intestinal clonotypes. The relevance of blood for diagnostic of tissue-directed TCR specificities (e.g. autoimmune), may thus vary with the tissue.
v. Very few of the clonotypes that expanded in tissues were found in the spleen and lymph nodes (LNs). However, dilution by large numbers of T cells from the broader polyclonal repertoire in SLOs would make tissue-specific clonotypes difficult to detect, and we cannot formally conclude that these cells are truly absent from SLOs, but they obviously constitute a much smaller fraction of the T cells

**Fig. 3.**
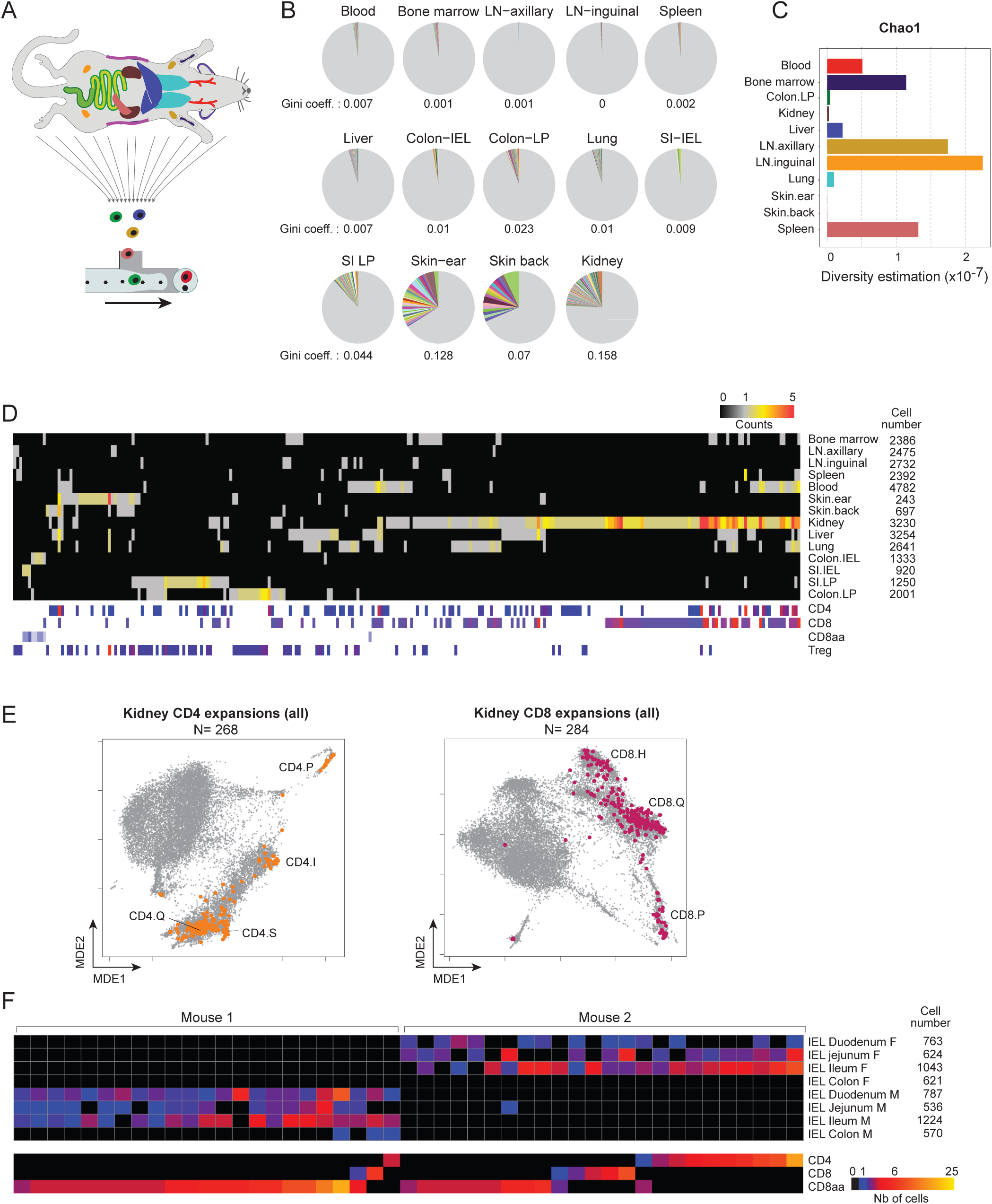
How the TCR repertoire distributes within one mouse. **A.** Many parenchymal organs and SLOs from a 7 week-old male B6 mouse were harvested, hashtagged and profiled together (combined IGT7/8/9). **B.** Pie chart of TCR clonotype distribution for the different organs (each downsampled to an equal 200 cells). The Gini index of diversity is shown for each organ. **C.** Chao1 estimate of the repertoire size in each organ. **D.** The organ of origin is shown for every clone detected at least twice in the whole mouse data, ordered by hierarchical clustering, with their cell-types shown below. Numbers on the right are the total number of full TCRαβ data in each organ. **E.** Phenotype of the cells expressing the kidney-preferential clonotypes is indicated by projecting them onto the MDE plots for CD4+ and CD8+ T cells (see Fig. 2D for reference). **F.** TCR clonotypes in IEL from different intestinal segments, from two different mice (one male, one female).

### Mechanics of TCR repertoire formation

Many studies have previously dissected the assembly of V, D and J elements to compose functional TCR units, but the depth and paired nature of the immgenT data warranted a re-examination of TCR rearrangement outcomes. In general, usage of *TRV* and *TRJ* segments fit with expectations (**Table S4A**). We also noted the well-described bias in *TRAV/TRAJ* (hereafter VaJa for short) pairing that derives from the preference for short-range rearrangements and the sequential opening of *TRAV* and *TRAJ* segments within the locus^36^ (**Fig. S3**). Overall, the numbers of nucleotides trimmed from the 3’ end of *TRV* and 5’ end of *TRJ* segments, and of N region additions (**Table S4B/C**) matched expectations from many past experiments. A large fraction of rearrangements lacked N nucleotides, even in immature thymocytes of adult thymi, refuting the misconception that the absence of N diversity implies a fetal/neonatal origin (perinatal T cells are indeed devoid of N addition^8^, but so are many T cells differentiated in adults).

Some TCRs differed from these overall norms, however. First, some had unusually short or long CDR3 loops (junctions of 9 AAs or less, 19 AAs or more). Very short junctions (in 968 out of 362,709 and 1,053 out of 497,192 *TRA* or *TRB* sequences, respectively) resulted from extensive trimming at the ends of *TRV* or *TRJ* segments, while very long junctions (251 and 554 instances, within *TRA* and *TRB* sequences), involved minimal DNA trimming and longer P and N nucleotide stretches, often retaining the complete *TRBD* segments in *TRB* (**Fig. 4A**). These loops of unusual length appeared independently in the two chains (**Fig. 4B)**. We found no strong association between extreme *TRA* and *TRB* junction length (long *TRA* junctions were not compensated by short *TRB* junctions, although there was a slight preference of long *TRB* junctions paired with shorter *TRA*; Chisq test, p = 0.66 and 0.002, respectively). This result suggests that the occurrence of extreme junction length in a given chain is sporadic and does not result from a genetic alteration of VDJ recombination in that cell. In these short and long junctions, we also observed a preference for different V and J genes in *TRA* and *TRB* (**Fig. S4A**). These disparately long CDR3 loops might generate lopsided pMHC/TCR interactions, and the longer loops might block interactions through steric hindrance. However, these extreme CDR3s do seem to interact productively with MHC molecules as they are found in all lineages (**Fig. S4B/C**), and there is precedent for structural plasticity in CDR3 loops that allows conserved docking onto pMHC^37^.

**Fig. 4.**
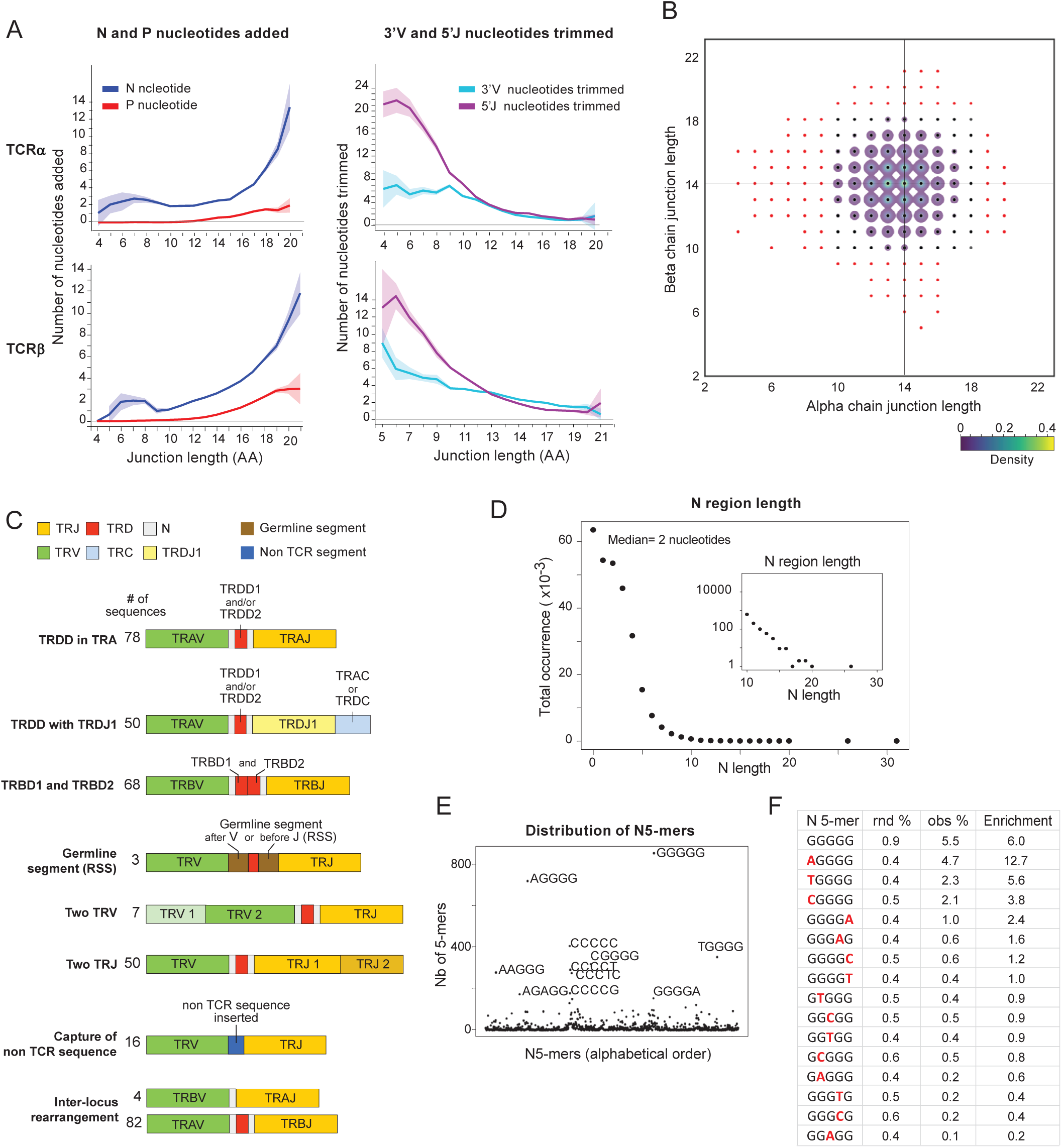
Molecular structures of VDJ joins. The entire non-transgenic immgenT dataset was analyzed for the presence of sequence structures and motifs at the recombination junctions. **A.** Averaged numbers of N and P nucleotides added (left) and of nucleotide trimming at the 3’ end of V and 5’ end of J segments (right) in *TRA* (top) and *TRB* (bottom) joins. The 95% confidence interval is shaded. **B.** Proportions of junctions of indicated length in *TRA* and *TRB* joins in the same cell. Red dots indicate unusually short or long junctions of nine AAs or less and 19 AAs or more, respectively. **C.** Schematic structure of non-canonical joins detected in the immgenT TCR dataset, color-coded as indicated **D.** Distribution of N nucleotides added per *TRA* join. **E.** Frequency of all possible N nucleotide 5mers. **F.** Table of G-dominant N nucleotide 5-mers, non-G bases in red. The expected frequency (rnd%) of each 5-mer, calculated from relative proportion of each base among mono-nucleotide N additions, is compared to the observed, “Enrichment” is the Observed/Expected ratio.

Some even more unusual patterns of rearrangements were observed (**Fig. 4C, Table S5**). These included TCRα and TCRβ transcripts with two V genes (**Table S5A-B**), or two J segments in tandem (**Table S5A-E**), likely involving cryptic recombination signal sequences (RSS). We observed a high number of TCRα sequences incorporating *TRD* segments, especially in γδ T cells (**Fig. S4B**), including sometimes rearrangements with *TRDJ1*. Most of these involved *TRAV*/*DV* genes that can be rearranged with either *TRAJ* or *TRDD* and *TRDJ* (e.g., *TRAV15-2/DV6-2*; **Table S5F**). This suggests that these joins derived from *TRD* rearrangements. We also observed *TRB* joins that included both *TRBD1* and *TRBD2* (**Table S5G**), and several instances of capture of genomic sequences from various unrelated chromosomes (13 to 50 nt; **Table S5H**). Finally, we observed joins in which germline sequences flanking *TRV*, *D* or *J* genes including their RSS were retained in the junction (**Table S5I**), and sequences resulting from trans-rearrangements between *TRA* and *TRB* genes (**Table S5J**). These “freak event” rearrangements were in frame and resulted in extremely long CDR3 loops. They were usually not accompanied by another normal rearrangement in the same cell, implying that they must partake in interaction with pMHC.

We also observed at very high frequency the previously described^38^ trans-splicing between the L-PART1 (IMGT nomenclature, encodes the 5’ untranslated and leader peptide) of *TRBV12-2* and the main V-EXON of *TRBV13-2*, accounting for 46,029 productive sequences (out of a total of 69,248 instances of transcripts first annotated as *TRBV13-*2 by V-QUEST; **Table S5K**). Similar trans-splicing involved members of the *TRBV12* and *TRBV13* families, albeit to lower extents, but few or zero for all other genes (**Table S5L**). Although the *TRBV12* and *13* families are intertwined on the genome, this preponderance cannot solely be linked to distance, and may involve a particular organization of the 3D nucleome that brings nascent RNAs from these genes into close proximity.

For an independent corroboration of these freak rearrangement events, we analyzed similarly a TCR dataset from mouse splenic T cells generated by split-pool technology, and made publicly available from Parse Biosciences ^39^. All of them were observed here as well (**Table S5M**), and at comparable frequencies, with the exception of the joins that include extra-locus elements (these may be filtered out by the Parse processing pipeline).

Much junctional diversity is created by random addition of nucleotides by terminal deoxynucleotidyl transferase (TdT) during the rearrangement process. The detailed call of N region nucleotides performed by the V-QUEST pipeline allowed a deep but conservative analysis of N-nucleotide additions (all junctions that might incorporate *TRD* segments or other freak structures were disregarded). N region assignments are often ambiguous for TCRβ due to D region remnants, so we focused on N additions of the VaJa joins (280,432 sequences, clonal amplifications collapsed). P-nucleotides were detected at low frequencies, averaging ∼0.4 in TCRα and ∼0.6 in TCRβ (**Table S4C**), and more frequent in longer junctions **(Fig. 4A)**. The general distribution of N additions (**Fig. 4D**) showed as expected a majority of joins with no or few N nucleotides (median=2 bases) but also with a non-negligible number of longer additions (1.5% with >7 bases, **Fig. 4D, inset**). For single-base additions, reflecting the known preferences of TdT, the expected ∼2-fold preference for G over other nucleotides was found^40^ (**Fig. S5A**). In keeping with earlier results^41^, the distribution of longer N additions was skewed towards G-rich homopolymers (**Fig. 4E)**, but the positions where other bases were found was clearly non-random (**Fig. S5B**, **Fig. 4F**): substitution of G by another base in the 5-mers was frequent at the first position (e.g. AGGGG), but not at internal positions (GGAGG was 50 times less frequent). The same applied to 6-mers. These biases are found in joins from pre-selection CD4+CD8+ double-positives (DP) in the thymus (**Fig. S5C**), thus unrelated to selection, rather a strong context-dependency of nucleotide preference of TdT.

### Recurring Rearrangements

TCRs used by MAIT and iNKT cells are very unusual^42^. They utilize predominantly a single VaJa combination, with a very limited joining diversity that results in a quasi-invariant CDR3. These canonical TCRα chains are paired with a more diverse, yet quite restricted, set of TCRβ chains. To ask whether these design principles are unique to iNKT and MAIT TCRs or shared more broadly, we interrogated immgenT data for such deviations (using a dataset of 299,000 paired TCRαβ sequences from all origins, with expanded clones collapsed). For each VaJa pair, we plotted the proportion of joins represented by the most frequent amino-acid junction against the total number of occurrences of the pair (**Fig. 5A**). VaJa pairs were widely distributed as expected^36^. The frequency of the most abundant TCRα junction ranged mostly between 4 and 8 %, but a group of outliers exceeded 20% for a fraction of pairs (hereafter AJ for “anomalous junction”; pval= 0.01 to 10^-12^ relative to the main distribution of junction frequencies per VaJa). These AJs included *TRAV11.TRAJ18* and *TRAV1.TRAJ33*, the prototypes of the category, but also ∼100 other VaJa pairs (**Fig. 5A**, 3 examples in **Fig. 5B**, listed in **Table S6A**). As is the case for MAIT and iNKT invariant alpha chains, the dominant junction in AJs was often complemented by very closely related junctions, differing by only one amino-acid (red in **Fig. 5B**). AJs were frequent events, ranging from 1/400 to 1/6000 of productive TCRα, far more frequent than their estimated probability of generation (OLGA Pgen 10^-5^ to 5x10^-8^). Indeed, the same number of VaJa joins generated probabilistically by the OLGA showed zero AJ, where the real data showed 35 with the same criteria (**Fig. S6A**). Strikingly, the distributions were completely different for TCRβ, where no VbJb pair yielded dominant junctions (**Fig. 5C**).

**Fig. 5.**
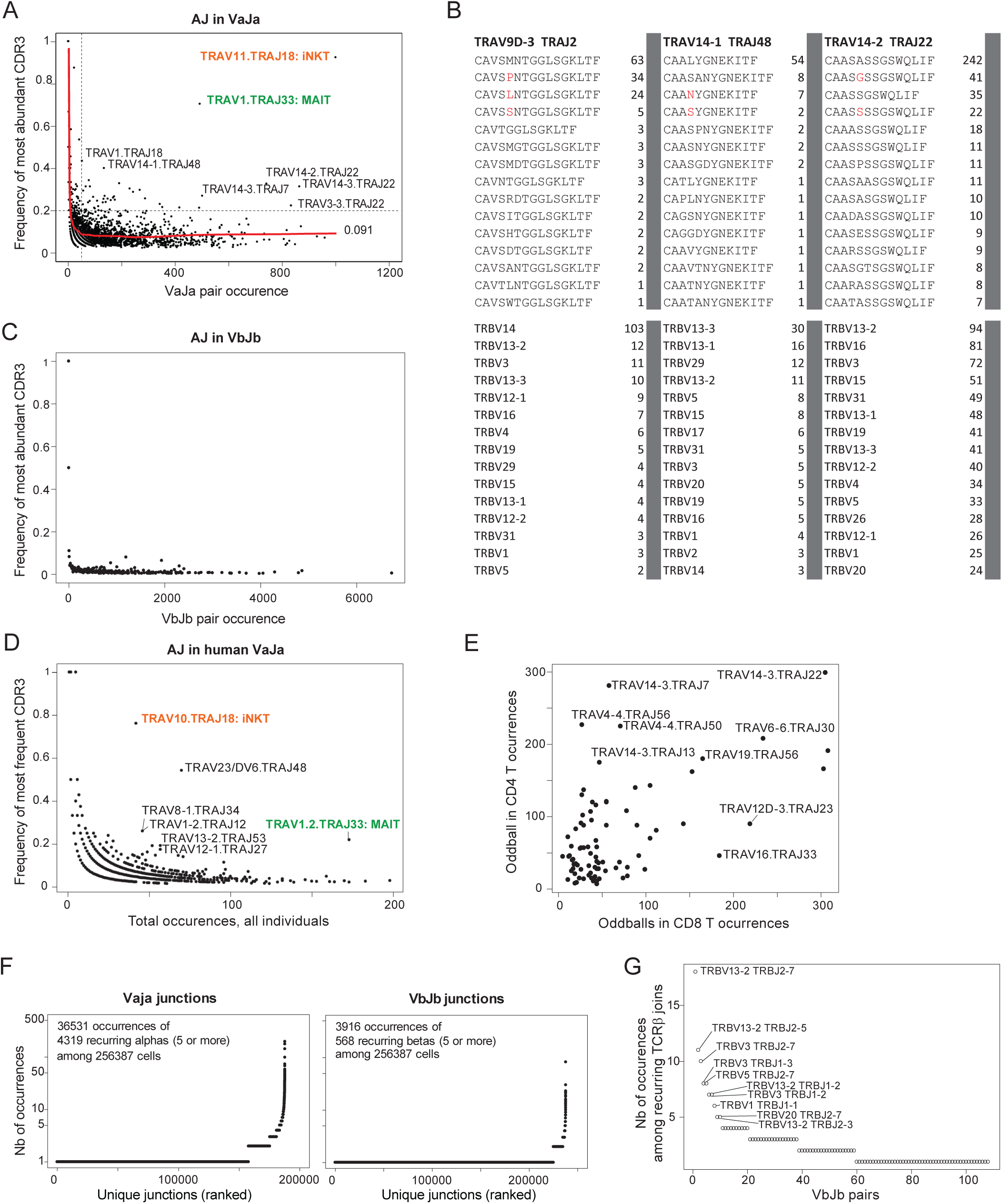
Recurrent rearrangements. **A.** For each possible VaJa pair, the total occurrence in the immgenT non-redundant dataset is plotted vs the frequency of the most frequent amino-acid junction encoded by the pair, the canonical VaJa pairs used by MAIT and iNKT cells are colored. The locally smoothed regression (supsmu function) between the two variables is shown in red. The dotted lines demarcate the outlier AJ pairs (occurrence>70 and main -junction-frequency >20%, ∼p<0.01). **B.** Three representative AJ pairs, with the sequence of their most frequent amino-acid junctions (top). When junctions differ from the dominant junction by only one amino-acid, the difference is shown in red. Bottom: Frequency of the *TRBV* segments most often associated with the same VaJa AJs and dominant junction. **C.** Same plot as in **A**, but for TCRβ. **D.** Same plot as in **A**, but for TCRα from human PBMC T cells, merged from 33 control samples. **E.** Differential frequency in CD8+ (x-axis) versus CD4+ (y-axis) T cells of mouse AJ VaJa joins, defined as shown in **A**. **F.** Count of the occurrence of unique VaJa junctions (left) and unique VbJb junctions (right). **G.** Count of the occurrence of VbJb pairs among recurring TCRβ joins.

Parse dataset from mouse splenic T cells by split-pool technology ^39^. Determining the frequency of the most abundant junction region for each VaJa pair again showed again the dominance of one junction for a subset of those pairs, involving the same VaJa pairs (**Fig. S6B**). Some *TRAVs* or *TRAJs* partake several times in AJ joins, but for any *TRAV* the junctional skew was only seen selectively: one to four of the VaJa pairs that use a given *TRAV* showed junctional skew, the others did not (**Fig. S6C**). Dominance by one CDR3 is thus a property of the VaJa pair, not of any *TRAV* or *TRAJ* individually.

Although not as frequent as iNKT and MAIT TCRαs, AJs are found between 0.1 and 0.01 percent of all T cells, collectively amounting to a sizeable fraction of the TCRα repertoire. Such frequencies were quite unexpected, relative to the generally accepted notion that any VJ pair is spread between a thousand or more CDR3 junctions. Most were present in spleen and SLOs, but some were enriched in the small intestine or in tumors (**Table S6A**). Invariant MAIT and iNKT TCRs utilize a restricted set of Vβ regions. This was also the case for several AJs (e.g. *TRAV9D-3.TRAJ2*) but not all (e.g. *TRAV14-2.TRAJ22*) (**Fig. 5B, Table S6A**). These data suggest that iNKT and MAIT TCRs are not isolated oddities, but only the tip of a larger iceberg of AJ TCRα joins with heavily biased distributions of CDR3 regions.

Interestingly, the same observation was made in a set of 55,000 human TCR sequences from blood T cells of 33 healthy donors (**Fig 5D**, **Table S6B**). Here again, MAIT and iNKT were the most skewed and abundant, but many other rearranged VaJa pairs also showed a dominant CDR3α, collectively representing 15 to 25% of the TCRs. These joins were not specific to any one donor, and were observed in ∼50% of the donors analyzed, even at this shallow coverage. Thus, AJs are not a quirk of mouse repertoires, but apply across species.

How do these dominant junctions originate? As was discussed for invariant iNKT and MAIT TCRs^43–46^, they could result from preferential rearrangement mechanics, repeatedly generating the same joins, or from strong selection which would pull out and expand cells whose TCRs carry the favored CDR3, or both.

With regard to rearrangement frequencies, several points are worth noting. First, most AJ joins are encoded by one dominant nucleotide sequence (**Fig. S6D, Table S6A, S6C**). Most are devoid of N nucleotide additions (mean= 0.45 N nucleotides vs 2.47 for whole data; **Fig. 4D**, **Table S6C**). This paucity might suggest preferential rearrangements driven by sequence homology at the V and J ends, as observed in TdT-deficient mice^47,48^. Closer examination of AJ joins revealed that they are complex, not as aligned as those from TdT-deficient cells, with variable nucleotide trimming on V or J sides. Many do include micro-homology, evidenced by the uncertainty on the origin of the corresponding bases **(Table S6C**). In early vertebrates like Zebrafish, sequence micro-homologies at fixed positions of germline V and J ends, possibly leftover from the primordial RAG transposon, strongly favor recurrent joins^49^. These features are lost in mammals, allowing greater diversification^49^. One might speculate that the recurrent AJ joins are remnants of the micro-homology control system, the recombination machinery still favoring micro-homologies, but now more diversely positioned in different VaJa pairs.

In terms of selection, it is important to stress that AJs do not result from clonal expansion, but represent repeated generations of related but distinct alpha-clonotypes. But their pairing with a limited set of beta-clonotypes, and the fact that different nucleotide sequences can contribute to the same alpha-clonotype (as for iNKT TCRs), suggest some selective pressure occurring after generation. The invariant TCRα chain of iNKT cells is thought to be very strongly selected, owing to its strong affinity for CD1d complexed with self-glycolipids, leading to a PLZF+ phenotype found mostly in the non-conventional group of cells (**Table S6A**). The other AJ TCRs were not dominantly represented in non-conventional T cells, but were mostly found in CD4+ and CD8+ T cells, with partial bias for one or the other (**Fig. 5E**). AJ distribution was better correlated between CD4+ T and Treg (**Fig. S6E**), presumably from common MHC-II restriction. This sharing of AJ TCRs by both MHC-I and MHC-II restricted lineages implies that there cannot be a single selective pressure, akin to ceramide+CD1d, driving their selection.

We also searched more directly for recurring rearrangements in the data, simply by counting the occurrence of single-chain clonotypes (nucleotide identity, after collapsing clonal expansions) that occurred repeatedly in different datasets (**Fig. 5F).** This hockey-stick plot showed a 10-fold larger number and incidence of recurrent clonotypes for TCRα than for TCRβ, as expected from the Chao1 estimates; indeed, many of the recurrent aN-clonotypes corresponded to AJ joins above (**Table S7A**). The recurrent bN-clonotypes stood out from the overall distribution, at markedly higher frequencies and uncorrelated to computational Pgen predictions **(Fig. S7A**, selected 298 occurring >5 times in **Table S7B**). Many were already present at comparable frequencies in pre-selection thymic DPs (**Fig. S7B),** suggesting recurring recombination events (all were associated with a different TCRα, so different from the public clonotypes described above). They were found in all lineages (**Table S7B**), but a special group was unique to CD8aa T cells (**Table S7C**): these correspond to the dominant clonotypes of CD8aa IELs (“Revere” in the terminology of ^50,51^, with exclusive usage of *TRBJ1-4* (**Fig. S7C**). Similarly, the recurrent aN-clonotypes also included a subset specific to CD8aa cells, with the previously reported Newbury clonotype, but also new clonotypes (**Table S7D**).

Setting these CD8aa-specific clonotypes aside, recurring bN-clonotypes involved a broad array of *TRBV* and *TRBJ* segments (**Fig. S7D**), but only a subset of VbJb pairs, with the dominance of only a few combinations (**Fig. 5G**), such as *TRBV13-2/TRBJ2-7*. It was also striking that, for this *V13-J2-7* combination, only a small minority of junction sequences were recurrent, all others being observed only once (**Fig. S7E**; this also applied to the other dominant combinations like *TRBV3-TRJ2-7*). No particular features explained these focused preferences (**Table S7E**): aside from having few N nucleotides, end-trimming was present but variable, some but not all included a micro-homology. Some included a clear D segment, which implies that the generation of these recurrent junctions required two independent recombination events.

Overall, the data reveal, mostly for TCRα but also to a lesser extent for TCRβ, recurrent patterns in the junction created by the recombination process. Unlike the clear micro-homologies between the termini of V and J genes that drive such phenomena in lower vertebrates^49^, the sequence grammar behind these preferences is unclear.

### Do unproductive rearrangements have any function?

Two thirds of TCR rearrangement events inevitably end up in out-of-frame unproductive joins, every cell having a second chance to rearrange productively its second TCR loci. Unproductive joins are thought to reflect the “raw” products of rearrangement mechanics, unaffected by any selection events. We thus analyzed the characteristics of unproductive joins in the immgenT dataset. Unproductive rearrangements of the second allele were detected less frequently than productive ones in the dataset (103,846 vs 849,127 altogether in the non-transgenic data), as expected since nonsense-mediated decay (NMD) lowers the abundance of untranslated mRNAs, resulting in less efficient capture during scTCRseq. V region usage showed the same frequencies as in productive joins. A totally unexpected result was observed when we measured *TRV* usage on the productive allele in cells that had an unproductive rearrangement on the other allele. As illustrated in **Fig. 6A** (**Table S8A-B**), the presence of an unproductive join in one cell diminished the proportion of the same *TRBV* in the productive join, by a factor of 3 to 10-fold depending on the *TRBV* (paired t.test p<10^-8^). This inhibition spread between members of a related family (*TRBV12* or *TRBV13*) suggesting that a sequence-or chromosomal location-dependent event was at play. The same phenomenon was observed for *TRAV* (p<10^-11^, **Fig. 6B**). We considered the possibility that this bias was an artefact in the TCR assembly algorithm, but several elements argued against this hypothesis. First, simulated raw reads from productive/unproductive pairs were correctly interpreted by CellRanger. Second, running the same data with the MiXCR suite^52^ resulted in the same distortions (**Fig. S8**). The most compelling argument for biological reality stemmed from a closer inspection of the interference between different chains; unproductive *TRBV13-2* joins decrease *13-2* and *13-3* joins, but also *TRBV12-2* joins. However, as shown above, transcripts annotated as *TRBV13-2* by V-QUEST include mRNAs transpliced from the leader of *TRBV12-2* onto the main exon of *TRBV13-2*. We thus parsed cells annotated as *TRBV13-2* into “true” *13-2* and those with the *12-2.13-2* chimera. The two molecules had completely different effects when unproductive, as only the unproductive joins involving the *12-2.13-2* chimera suppressed productive *12-2* rearrangements, while the unproductive “true” *13-2* transcripts did not (**Fig. 6C**). This split indicated that the suppression phenomenon was tied to specific sequences at the 5’ end of the rearranged TCRβ mRNA. We tentatively interpret these results as indicating that a failed unproductive TCR transcript is not inert, but functions as a long non-coding RNA (lnc). Lncs regulate chromatin structure and transcriptional activity in a sequence specific fashion^53^, and an unproductive TCR mRNA could specifically repress in *cis* the accessibility of its own promoter region and also inhibit in *trans* the accessibility of its homolog for rearrangement, perhaps by promoting the assembly of repressive nuclear condensates in a heterochromatic neighborhood. There is no known example in biology where an mRNA carrying a nonsense mutation on one allele inhibits the expression of the other allele (which would be counter-productive). The opposite has been documented, decay of a nonsense mRNA leading to the compensatory upregulation of its functional homolog^54^. However, immune receptor genes are different in that they naturally produce many nonsense genes. A mechanism that specifically silenced unproductive *TRV* promoters could be beneficial, avoiding wasteful transcription and relieving the overload on NMD mechanisms. Losing the ability to rearrange the homologous *TRV* would be un-consequential, as many other *TRVs* would remain available. Another interpretation of these observations is the reverse: a translated in-frame TCR mRNA might suppress the expression of a matched unproductive transcript, lowering the odds of its assembly and detection in the single-cell data. This alternative mechanism would also rely on close homology between the silenced promoter and the mRNA 5’ end.

**Fig. 6.**
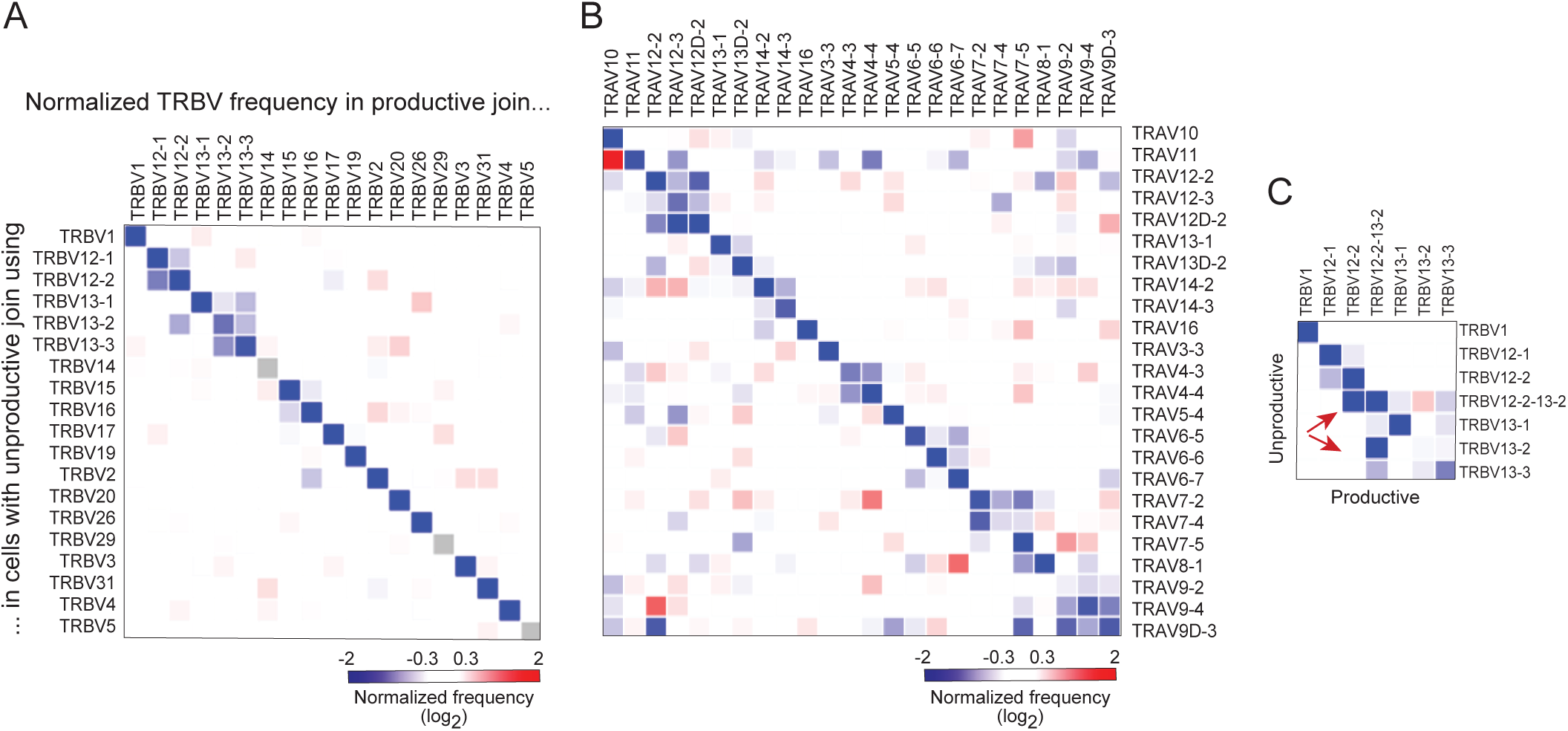
Interference between unproductive and productive joins. **A.** The proportions of productive/unproductive pairs of rearranged *TRBV* in the primary and secondary alleles of each cell was counted, across the non-redundant non-transgenic dataset. For clarity, values are normalized vs the mean usage of each *TRBV*. **B.** As in A, for unproductive/productive pairs of *TRAV*. **C.** Transcripts initially assigned to *TRBV13-2* by V-QUEST were re-analyzed and separated into those that use the trans-spliced transcript (PART1 leader of *TRBV12-2* spliced onto main exon of *TRBV13-2*) or the “pure” *TRBV13-2* transcripts, and the productive/unproductive pairs were counted as in **A**, Arrows demote that productive *TRBV12-2* usage is under-represented in cells with a unproductive trans-spliced variant, but not by the unproductive “pure” *TRBV13-2*.

### Allelic exclusion

Allelic exclusion in TCRs, in the textbook definition, ensures that each T cell expresses a single TCR with one binding specificity. It is enabled by the sequential and controlled rearrangement of *TCR* genes during thymic differentiation. The β-chain locus rearranges first. If successful, the TCRβ chain pairs with a pre-Tα chain to form the pre-TCR complex. If the pre-TCR complex folds effectively and reaches the cell surface, it signals and inhibits rearrangement of the second TCRβ allele. After further differentiation and expansion, the TCRα locus opens and rearrangement occurs. Allelic exclusion for TCRα is thought to be leakier^55–57^, either because signals from positive selection are less effective than those from the pre-TCR at stopping rearrangement, or because α-chain rearrangement can occur on both alleles at once. Physiological or pathological roles for these dual-α T cells have been proposed, by enhancing cross-reactivity or autoimmune potential^56,58–60^.

Some exceptions to absolute exclusion at the TCRβ locus have been reported through the years^61–65^, and we revisited these notions in the particular cell populations of the immgenT dataset. Of 228,373 cells with both productive TCRα and TCRβ rearrangements, 23,170 had two productive α-chains (10.14%), and 13,082 had two productive β-chains (5.72%), suggesting less difference in allelic exclusion than expected at the mRNA level. These did not appear to derive from doublets, as the read distribution was similar for dual-α, dual-β and single-chain cells (**Fig. S9A**; cells with dual TCRα and TCRβ did seem enriched for doublets). These dual chain cells were represented quite uniformly across different lineages, around 9-10% and 5-6% for dual-α and dual-β, respectively (**Fig. 7A**), and not enriched in Treg cells as had been suggested. Single-cell profiling by microfluidic encapsulation can suffer from contamination from ambient RNA. As an independent verification, we analyzed the split-pool dataset of mouse spleen T cells mentioned above ^39^. Of 122,735 cells with both TCRα and TCRβ, 7.1% showed a second productive TCRα join, and 6.2% a second TCRβ join, in good accordance with the immgenT dataset (the overall proportions are lower in the Parse data because of a slightly lower discovery rate).

**Fig. 7.**
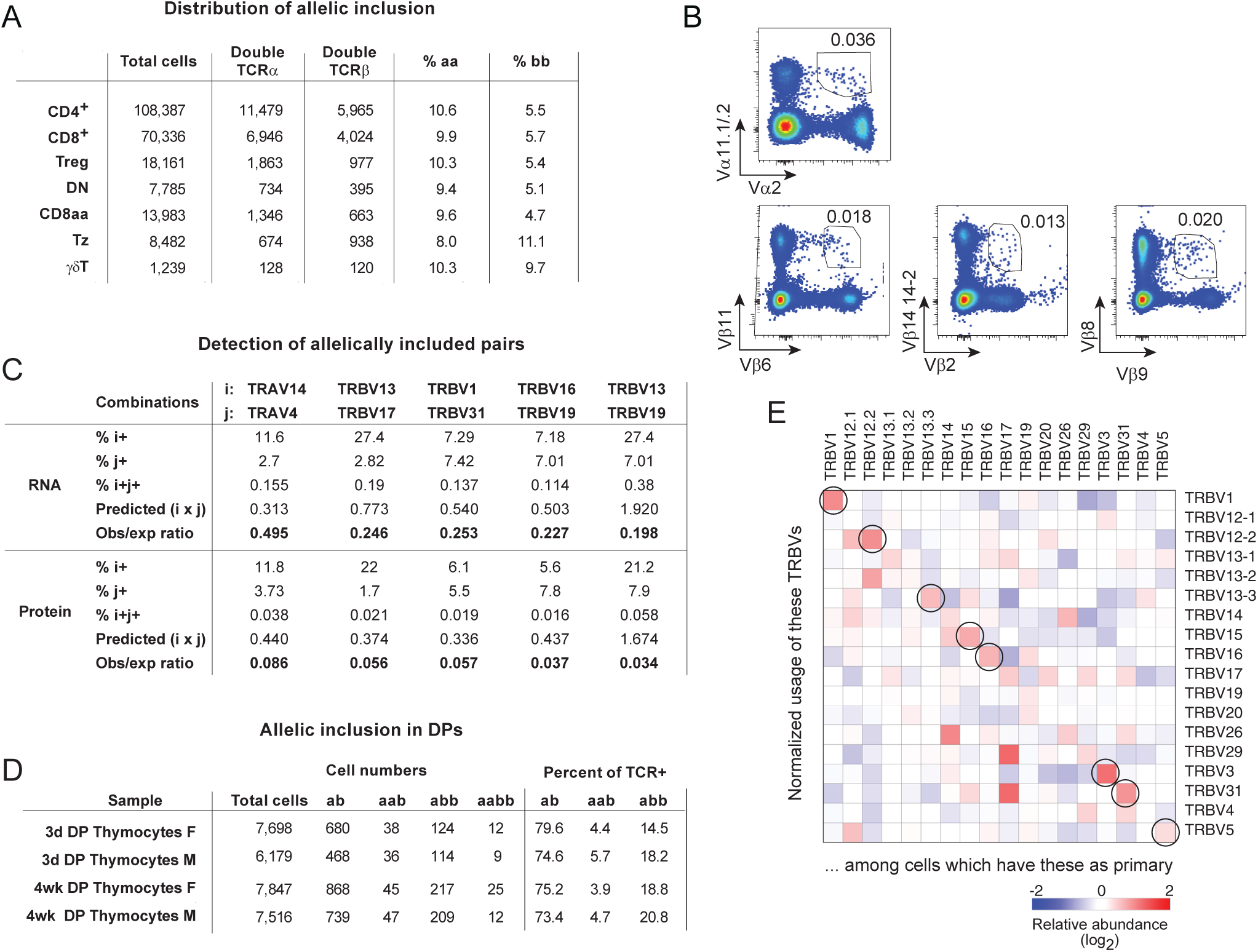
Leaky allelic exclusion at both TCRα and TCRβ. **A.** For all major immgenT lineages, number and percentage of cells that have two productive TCRα or TCRβ chains and at least one alpha and one beta (peripheral T cells only). **B.** Validation by flow cytometry of dual-positive cells that display the products of two *TRAV* (top) or two *TRBV* (bottom) genes. Classic names of V regions detected by the antibodies are shown (*Vα2* = *TRAV14*, *Vα11.1/.2* = *TRAV4-4/DV10*, *Vβ6* = *TRBV19*, *Vβ11* = *TRBV16*, *Vβ2* = *TRBV1*, *Vβ14* = *TRBV31*, *Vβ9* = *TRBV17*, *Vβ8* = *TRBV13*). **C.** Comparison of detection of single or dual expressor cells at the RNA and protein level, for the *TRBV* or *TRAV* combinations shown (i, j indexing). RNA is from immgenT TCRseq, protein from flow cytometry (as in **B**). “Predicted” double expressors is the product of the proportions of both V regions, were there no counter-selection. “Observed ratio” is the ratio of this predicted value over the proportion of i+j+ cells actually detected. **D.** Numbers and proportions of single and dual expressor cells among immature DP thymocytes from young mice. The lower proportion of aab cells compared to the periphery (**A**) is expected since only a minority of DPs have rearranged TCRα chains. **E.** Proportion of cells with a productive *TRBV* on the primary TCR (beta in SummaryTable) combined with the same or another productive *TRBV* join on the second allele (beta2 in SummaryTable). Normalized vs the mean frequency of each *TRBV* among productive joins.

Flow cytometry staining verified that these dual-β cells do exist and were not scRNAseq technical artefacts (**Fig. 7B**). We further confirmed this staining by performing TCRseq after sorting Vβ6Vβ11 dual-positive cells, which indeed contained two productive TCRβ mRNAs (**Fig. S9B**). Dual-α cells were much easier to detect, in line with prior reports^56,61,63,64^. Compared to the proportion expected by combining the individual frequencies of each member of a pair, the observed proportions of dual-positive cells were somewhat reduced (2 to 5-fold) in the scRNAseq data, but much more so in the flow cytometry (12 to 30-fold) (**Fig. 7C**). The simplest interpretation is that there is competition at αβ pairing, and that T cells go more effectively through positive selection and homeostatic maintenance if they express one dominant TCR.

In data from thymic immature CD4+CD8+ (DPs) cells, dual-β cells were already present (**Fig. 7D**). These results indicate that dual-β cells are generated early, during normal TCRβ rearrangement, and not because of secondary editing in mature T cells. Overall, the frequency distribution of *TRBV* in the first and second chains was quite comparable in the primary and secondary rearrangements of these dual-TCR cells (**Fig. S9C**). We asked whether the combinations of V regions used by dual-β cells were random. This proved not to be the case, with a preference for the same *TRBV* being employed twice in the same cell (**Fig. 7E**). For instance, a cell expressing *TRBV1* as its first β-chain was 3.8 times more likely than by chance to utilize *TRBV1* for its second rearrangement. This bias was true for 8 of 21 *TRBVs* (**Fig. 7E**). Although the TRBV locus does not display the same ordered opening as *TRAV*, it might be that the loop extrusion and RAG-scanning process preferentially target the same *TRBV* in a given proT cell.

### Order of rearrangement

Another textbook notion is the TCRβ>TCRα order of rearrangements, and the cell divisions between TCRβ and TCRα rearrangements, which predict that groups of T cells can carry the same TCRβ clonotype, associated with different TCRα that rearranged in different descendants of the same β-rearranged cell. The reverse, one TCRα associated with several different TCRβ, should be absent. We tested this notion by identifying occurrences of one TCRβ clonotype (full nucleotide identity) associated with different TCRα, and *vice versa*. Importantly, this search was done iteratively within each immgenT sample, not across samples, to identify repeats happening within one mouse, not public TCRs. We identified 1,724 instances of the same TCRβ clonotype paired with different TCRα, and a surprising 1,132 instances of the same TCRα paired with different TCRβ clonotypes (hereafter “1a2b”, **Table S9**). The latter did not seem artefactual, since they had a normal distribution of sequencing reads and UMIs for each contig, were not doublets (**Fig. S10A**), had the usual levels of end trimming and N additions, and were generally represented across all lineages (**Fig. S10B**). Artefactual contamination by ambient RNA during was also ruled out for most 1a2b events (detailed in legend to **Fig. S10C/D**). Alpha chains in 1a2b pairs had a broad and normal range of Pgen probabilities (mean= 4.4x10-7 vs 3.9x10-7 for the whole data), and only six of 1,132 1a2b situations involved the frequent TCRα joins described above (**Table S9**), suggesting that they did not simply stem from recurrence of favored TCRα rearrangements.

A possible interpretation of 1a2b events is that TCRα rearrangement and expansion can happen before TCRβ in some pro-T cells. Accordingly, we searched for cells with only TCRα rearrangements among immature thymocytes. The matched UMAPs computed from RNA and surface-protein information (**Fig. 8A/B**) in combined IGT17/18 datasets (normal B6 thymus, double-negatives (DN) oversampled, analyses performed on CD73-negative cells to exclude recirculation) illustrate the value of this combined information served on the immgenT Rosetta viewer (detailed in legend to **Fig. S11A/B**). Two regions of RAG expression were readily identified, in early DN and immature DP (**Fig. 8C**). These RAG+ clusters contained cells with only TCRα or TCRβ rearranged (**Fig. 8D**). The more abundant β-only DNs were expected, since this is where RAG expression and TCR rearrangement starts, but α-only cells were unexpected, if TCRα rearrangement only occurs at the later DP stage. These α-only DN cells did not result from dropout of TCRβ transcripts in TCRα+TCRβ+ cells (as happens in the single-positive clusters) because there were few TCRα+TCRβ+ cells in the DN clusters (**Fig. 8E**), They did not result from ambient RNA, as their proportions (5.7% of cells in cluster-0) were much higher than ambient RNA levels (∼0.3% of cells for IGT17, Supplementary Note), and TCRs of α-only DNs were not seen in other cells in the sample (including two pairs of duplicated TCRα; **Table S9B**). Using serial gating in Rosetta, including filtering out of CD73+ recirculating cells (**Fig. S11C**), we displayed CITEseq data as the classic CD25/CD44 plot to parse DN subsets (**Fig. 8F**).Interestingly, α-only DN cells were restricted to the DN2/DN3 compartment, and progressed very little to the DNA cycling clusters (**Fig. 8G.H**), in contrast to β-only cells which progressed to DN4 and to cell cycling (**Fig. 8I,J**). DNs with both chains mostly matured and entered the DN4 cycling cluster (**Fig. 8K,L**). We interpret these results as indicating that both TCR loci can rearrange in RAG-positive DN cells. The TCRβ locus does so more effectively, and pairing with the preTCRα chain rapidly drives the cells to proliferation. TCRα does not have this option, and α-only DNs remain at DN3 until a productive TCRβ rearrangement occurs and the TCRαβ pair enables progression and active division starts. Although not exclusive, the more efficient early rearrangement at TCRβ, together with the amplification provided by the preTCRα chain, explains its dominance and why early TCRα rearrangements may have gone un-reported. We speculate that the concomitant rearrangement of both TCR chains in DNs represents an evolutionarily early solution, before preTCRα allowed amplification of early TCRβ-expressing clones. These interpretations are consistent with prior observations of sizeable T cell populations in *Ptcra*-deficient mice and patients^66,67^, although they may not suffice to explain 1a2b cells.

**Fig. 8.**
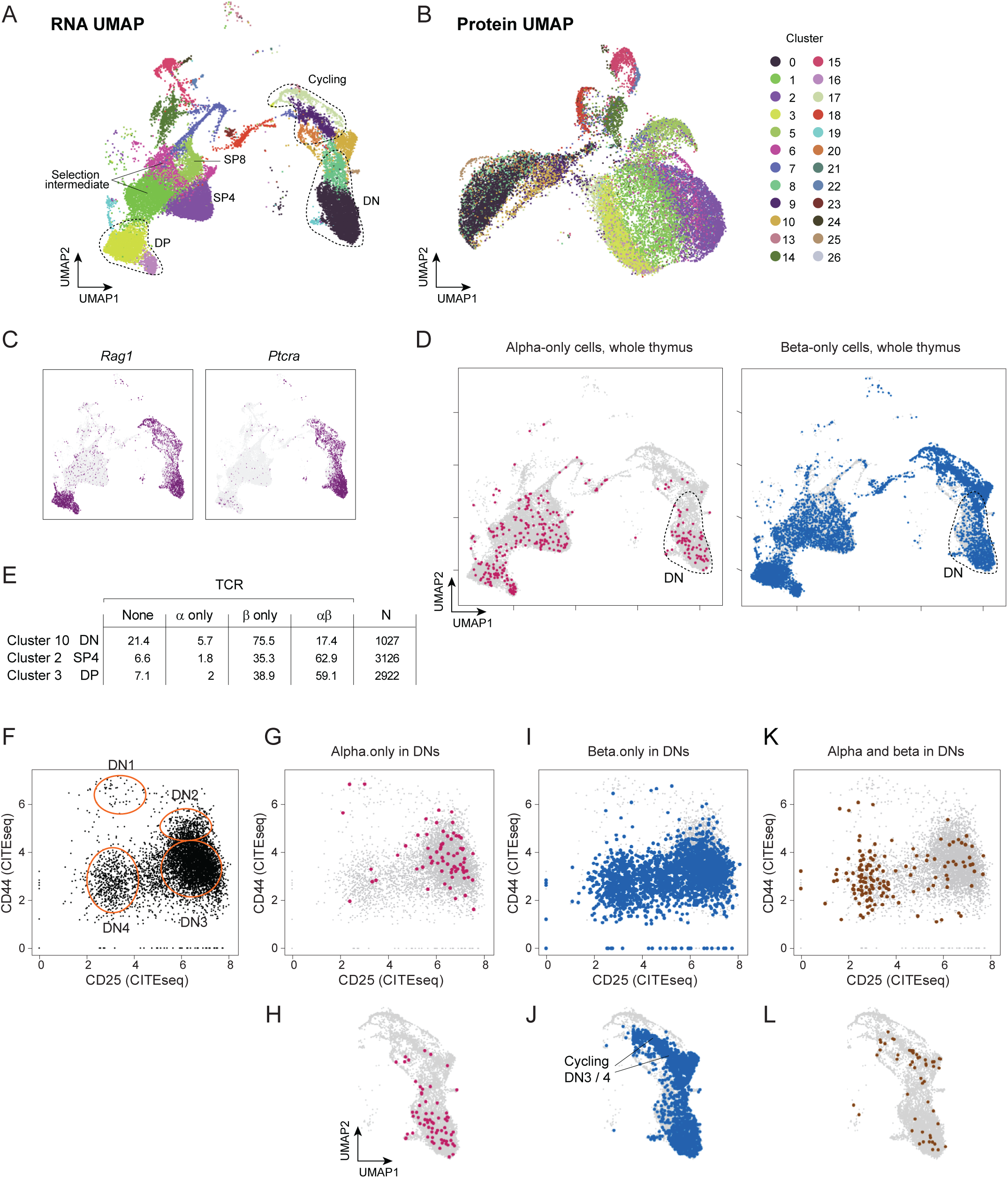
Immature DN thymocytes rearrange TCRα as well as TCRβ. **A.** UMAP representation of thymocytes from gene expression in the combined IGT17/18 datasets (total thymocytes, but downsampled to avoid an excess of DPs, and suspect dying-cell clusters filtered out). Color-coded according to Seurat clusters. Annotation derived from expression of keymarker genes (see **Fig. S11A/B**). **B.** Same data as A, but UMAP representation derived from the cell surface marker data. Color-coding according to clusters in the RNA UMAP in A. **C.** Expression of *Rag1* and *Ptcra* gene, same UMAP as in A. **D:** Thymocytes with only TCRα rearrangements (left) or only TCRβ (right) in the same UMAP as in A. **E.** Proportion of cells with TCRα rearrangements only, TCRβ only, or both in DN, SP4 and DP clusters from A. **F.** CD25/CD44 profile (from CITEseq data) of DN cells (gated as CD73-TCRb-CD4-CD8-). The classic DN subsets are circled. **G, I, K**: Same plot of DN cells as F, but showing the cells with only TCRα rearranged, only TCRβ, or both. **H, J, L**. Same cells as in G/I/K, but shown on the DN section of the UMAP in A. The position of cycling DN3/4 derives from **Fig. S10B**.

### Germline selection molds the TCR

After positive selection in the thymus, TCR clonotypes expressed by mature CD4+ and CD8+ T cells are restricted to interacting with MHC-I or MHC-II molecules, respectively. Although this partition is not absolute, it does constitute a major fault line for T cells, segregating the type of antigen recognized with the cells’ effector functions. This dual ability of TCRs raises an evolutionary conundrum: how can TCR components be evolutionarily selected for the ability to interact with two different ligands, which are themselves highly variable between individuals of a species^68–72^? It has long been known that individual Va and Vb regions are preferentially utilized by CD4+ or CD8+ T cells ^73,74^, through preferential contacts that involve CDR1 and CDR2 loops, and are further modulated by CDR3 ^75,76^. In the immgenT data, every V region was observed in both MHC-I and MHC-II restricted cells, confirming that each is potentially compatible with both MHC-I and MHC-II molecules. In keeping with classic mAb-based observations, however, there were marked differences in usage of several *TRAV* and *TRBV* genes between CD4+ and CD8+ T cells (**Fig. 9A, Fig. S12A, B, Table S10A-B**), but far less between MHC-II restricted CD4+ and Treg cells, or between MHC-I restricted CD8+ and CD8aa populations. These biases were mostly independent of the J regions (**Fig. S12C, D**) which showed only slight preference for CD4+ or CD8+ T cells. The differential selection of V regions into CD4+ or CD8+ pools could theoretically reflect stronger TCR/MHC interactions leading to enhanced representation after positive selection, or conversely weak interactions that diminish the probability of selection. Comparing *TRBV* usage in pre-selection thymic DPs vs mature CD4+ and CD8+ T cells showed that both are true: for instance *TRBV12-1* increases in frequency in mature CD8+ (**Fig. 9B**), and decreases in mature CD4+ (**Fig. 9C**), relative to DPs (and the reverse for CD4-biased *TRBV20* or *31*). Thus, evolutionary selection allows interactions of most V regions with both MHC-I and MHC-II, but for some of them do preferentially interact with one of the two classes, at the detriment of the other.

**Fig. 9.**
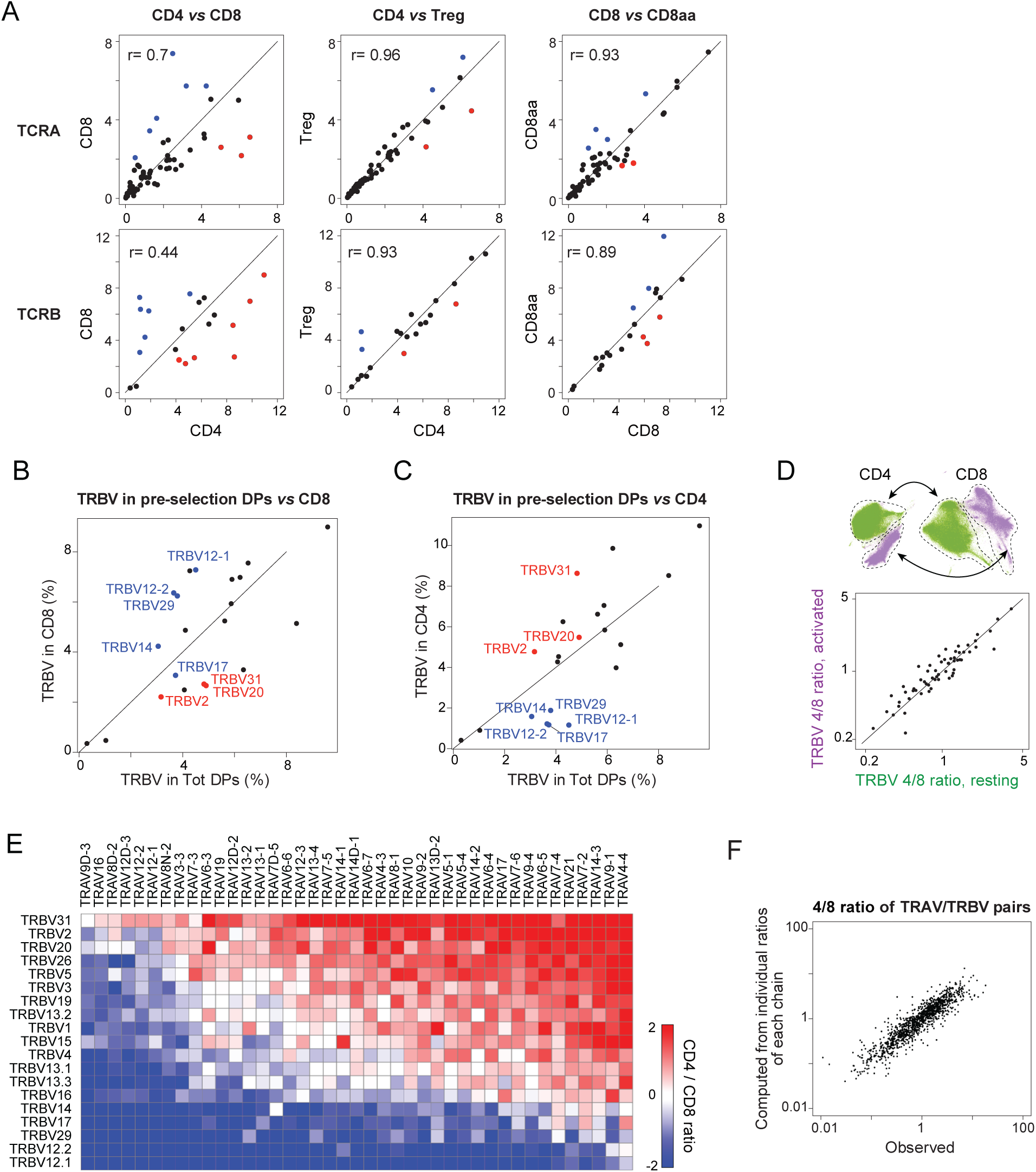
Preferential V region usage after positive selection on MHC-I or MHC-II molecules. **A.** Comparison of *TRAV* (top) or *TRBV* (bottom) frequencies among productive joins, for the indicated sets of lineages. Red and blue dots are *V* genes that show statistically significant differential usage between CD4+ and CD8+ lineages (left panels). **B.** Comparison of *TRBV* frequencies in immature pre-selection DP (x-axis) vs mature CD8+ T cells. Red and blue color-coding as defined in **A**. **C.** Comparison of *TRBV* frequencies in immature pre-selection DP (x-axis) vs mature CD4+ T cells. Red and blue color-coding as defined in **A**. **D.** Ratio of *TRBV* usage frequency in CD4+ vs CD8+ T cells, calculated for naïve/resting (x-axis) or activated/experienced (y-axis) clusters defined at top. **E.** Ratio of usage in mature CD4+ vs CD8+ T cells for all *TRAV/TRBV* pairs (for robustness, selected for individual *V* gene usage >2%). **F.** As in **E**, ratio of usage of every *TRAV/TRBV* pair in mature CD4+ vs CD8+ T cells (x-axis) plotted against the product of the CD4/CD8 ratios of each *TRAV* and *TRBV* constituting the pair.

In theory, V region usage biases could result from preferential selection in the thymus, or from TCR/MHC interactions in the periphery, as trophic signaling or as activation signals. We hypothesized that the latter would be manifest as different CD4/CD8 ratios in cells categorized as naïve and as antigen-experienced in the immgenT gene expression data. This was not the case, and CD4/CD8 biases were essentially identical in the naïve-to-naïve and activated-to-activated comparisons (**Fig. 9D**), suggesting that they are independent of TCR/MHC contacts during T cell activation.

Recognition of pMHC complexes depends on contributions from both TCR chains, but does the propensity towards MHC-I or MHC-II interactions simply sum the bias of both chains, or are there unexpected combinatorial effects? The CD4/CD8 ratio of usage frequencies for all α/β pairs (**Fig. 9E**) showed that the bias exhibited by a given pair largely sums the effect of the two components, with perhaps a slight dominance of the TCRβ chain (*TRBV12* predominant in CD8+ irrespective of *TRAV*, similarly for *TRBV31* in CD4+). Indeed, simply averaging for each VαVβ pair the frequency in each lineage of its two constituents yields a CD4/CD8 bias very closely related to the observed values (**Fig. 9F**).

Further complicating the germline selection conundrum is that MHC-I and MHC-II restriction often involves several molecules, in mice MHC-I H2-K and H2-D vs MHC-II H2-A and H2-E (hereafter A and E). We crossed transgenic and knockout mice to generate mice expressing only A or E, and asked how V region usage compares in CD4+ cells selected on A or E molecules only. *TRAV* usage showed modest biases in E- vs A-restricted CD4+ T cells, mostly 2-fold or less, which proved significant upon permutation testing (**Fig. 10A, Table S11A**). Vα regions preferentially used in CD8+ T cells were equally under-represented in CD4+ T cells from E-only mice (**Fig. 10A**), implying that MHC-I versus MHC-II bias operates similarly for recognition of A and E.

**Fig. 10.**
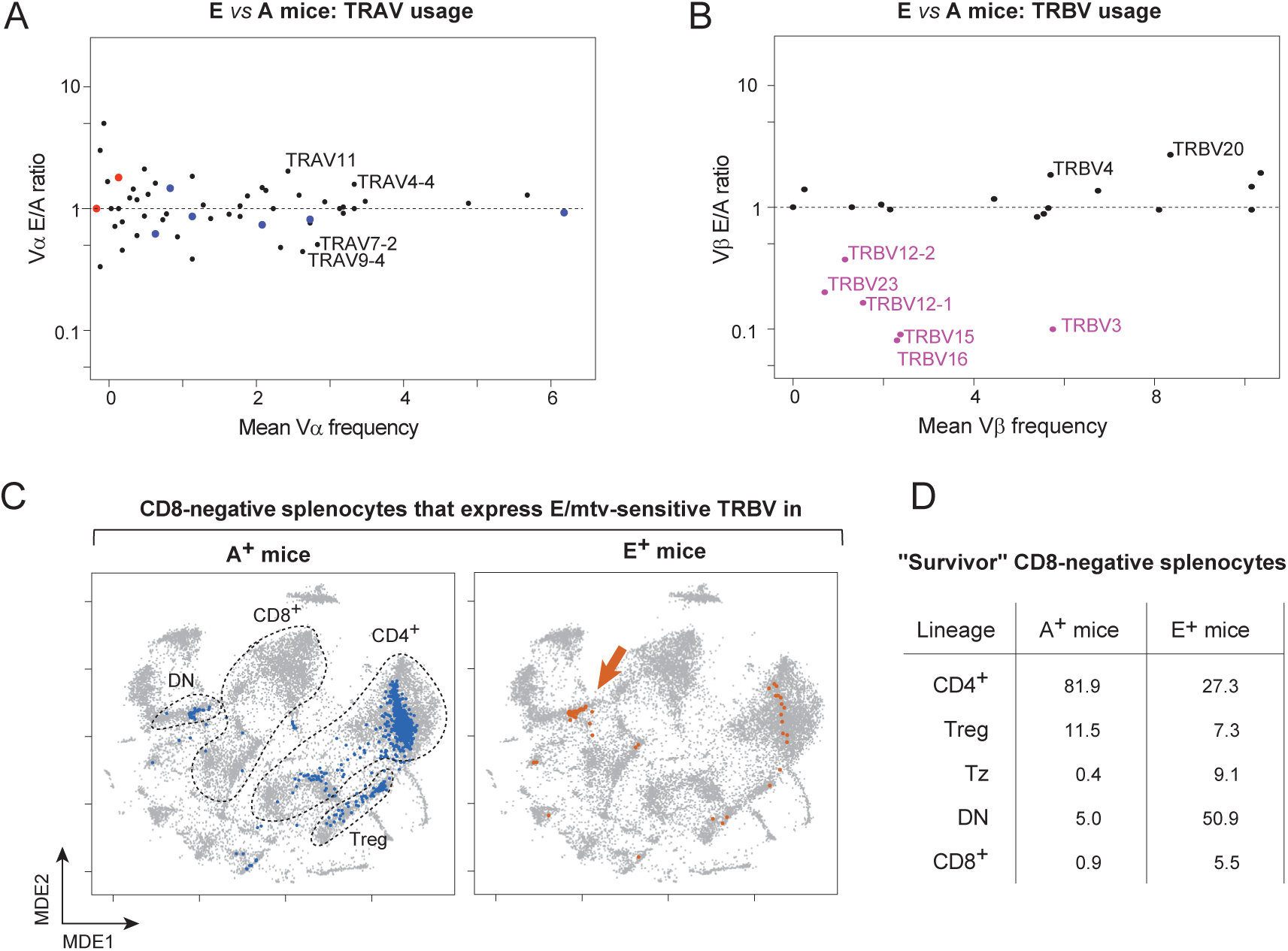
Repertoire selection by H2-A or H2-E MHC-II molecules. Mice expressing only H2-E (aka I-E) MHC-II molecules were bred by intercrossing the A-deficient Ab° line with the Eα16 transgenic, which complement genetically deficient Eα expression of B6 mice. B6 mice naturally express only the H2-A molecule. **A.** Comparison of *TRAV* usage in CD4+ splenocytes (IGT102/103) from E-only and A-only mice. Red and blue dots indicate the *TRAVs* differentially represented in CD4+ vs CD8+ T cells of B6 mice, as defined in **Fig. 9A**. **B.** Comparison of *TRBV* usage, as in A. Purple letters denote *TRBV* genes whose usage declines sharply in E+ mice owing to deletion by endogenous *Mtv*-encoded superantigens (*TRBV12* and *TRBV23*, *TRBV15* and *TRBV16* correspond to the Vβ5 family, Vβ11 and Vβ12, respectively). **C.** The position of CD8-splenocytes that express *Mtv*-sensitive *TRBVs* (per panel **B**) is projected onto the immgenT “all-T” MDE plot, for cells from A+ or E+ mice. The shift to DN population in “deletion survivors” of E+ mice is indicated by an arrow. **D.** Quantitation of the results from **C**, distribution of CD8-splenocytes within the main immgenT lineages.

On the other hand, some more dramatic under-representations of Vβ regions in E-positive mice were observed (**Fig. 10B, Table S11B**). These corresponded exactly to the classic clonal deletion induced by Vβ-specific endogenous superantigens (*Mtv8, 9* and *17)* present in the B6 genome^77,78^. We then interrogated the gene expression data to ask what phenotype is adopted by “survivors” of clonal deletion, i.e. splenic T cells that use those *TRBVs* in E+ mice. No deviation towards Tregs or “anergic-like” CD4+ phenotypes was observed. Instead, there was a marked increase in CD4-CD8-DN cells, mostly of subclusters DN-cl29 and 39 (**Fig. 10C/D**). These observations suggest that the escape from sAg-induced negative selection does not merely involve the avoidance of signaling-induced cell death, but differentiation to an alternative fate, described in more detail in an accompanying manuscript (**immgenT-Tz**). Collectively, these observations imply that the evolutionary pressures that ensure matching TCR elements to A operate similarly for E (even though A and E have only ∼50% sequence identity), with no “specialization” of the V regions. They also bring back the old E puzzle: although the A/E duplication is evolutionarily conserved (HLA-DR/HLA-DQ in humans, and present in all placental mammals), why do mice continuously attempt to eliminate E function, either through genetic mutation^79^ or immunologic inactivation of T cells that would interact with it?

## DISCUSSION

TCR repertoire studies in the mouse have a long history, but the immgenT context provided an unprecedented perspective on the paired αβTCR repertoire, not simply in blood and lymphoid organs but in many tissues, and in direct relation to information on lineages and cell phenotypes. Compared to traditional cytometry studies limited by mAb availability, the data could query the behavior of all genes through selection, and the single-cell framework avoided the issues with one-chain massively amplified approaches. The results bring a number of unexpected observations, and should open to fertile experimentation. Many of the observations are discussed in the sections above; we will not restate them here, but some overarching points may be worth stressing.

It has been known for some time that the TCR repertoire is lumpy, with a great range of clonotype frequencies^5,11,17,28,76^. The nature of the dataset allowed us to parse what portion is due to clonal expansion, highlighting a diversity of mechanisms that underlie this heterogeneity. Some clearly reflect immunological selection, like the public clonotypes in *M. tb* or SFB infections. Others suggest non-random mechanisms during recombination, perhaps evolutionary shortcuts to recognize common antigens, or to favor matches to MHC: the AJ TCRα joins, the less frequent but equally puzzling recurrent TCRβ joins, the Revere family of TCRs used by CD8aa T cells. We could not detect uniform sequence features, like those present in TCR genes of lower vertebrates^49^, that would explain the fine skew in recombination output. One might speculate that a more complex code evolved at the ends of mammalian TCR genes, or some form of templated resealing of RAG-created joins.

Although the partitioning of T cell clones between tissues is an established concept, we were surprised how highly compartmentalized the TCR clonotypes appeared, with limited overlap between tissues, implying strong effects of local antigen exposure and/or migration patterns. The blood repertoire showed limited overlap with tissues clones. Whether this reflects limited circulation of T cell clones amplified in tissues or dilution by the larger SLO repertoire (likely both), the practical implication is that blood may be a poor proxy for tissue-specific repertoires and immune responses.

Finally, the results suggest that TCR germline regions are evolutionary adapted to recognize different classes of MHC molecules with a soft combinatorial code, rather than being strictly specialized for MHC-I or MHC-II. But this structural matching remains able to cope with wild outliers in CDR3 loop lengths.

## ACKNOWLEDGEMENTS

We thank Drs. D. Schatz, T. Boehm, M. Krangel, C. Bassing, P. Marrack, M, Flajnik, B. Malissen, and K.C. Garcia for insightful discussions; K. Hattori, C. Araneo and the Klarman Observatory for help with mice, cell sorting and single-cell profiling. We are grateful to Parse Biotechnology for making public their dataset of mouse T cells. This work was funded by a grant from the NIH to the ImmGen consortium (R24-072073), by funds to IMGT from the Scientific Research National Center (CNRS), the Université de Montpellier, the Institut Universitaire de France (IUF) and Institut du Développement et des Resources en Informatique Scientifique (IDRIS) under the allocation 036029 (2010–2026) made by GENCI (GrandEquipement National de Calcul Intensif).

## AUTHOR CONTRIBUTIONS

IM and VP performed the experiments, MC, LY, SC, OC, BV, and CB the computational analyses; DZ and CB designed the overall study; VG, SK, DZ and CB provided funding and oversaw the experiments; MC, SC, and CB wrote the manuscript with input from all authors.

## COLLABORATORS

Participants in the immgenT Project include:

Aaron Liu^1^, Alexander Chervonsky^2^, Alexandra Cassano^2^, Alia Welsh^3^, Amir Ferry^11^, Ananda Goldrath^11^, Andrea Lebron-Figueroa^5^, Ankit Malik^2^, Anna-Maria Globig^4^, Antoine Freuchet^2^, Bana Jabri^2^, Charlotte Imianowski^6^, Christophe Benoist^5^, Claire Thefaine^7^, Dan Kaplan^6^, Dania Mallah^5^, Dario Vignali^6^, David Sinclair^5^, David Zemmour^2^, Derek Bangs^8^, Domenic Abbondanza^2^, Enxhi Ferraj^9^, Eric Weiss^6^, Erin Lucas^7^, Evelyn Chang^9^, Gavyn Chern Wei Bee^10^, Giovanni Galletti^11^, Ian Magill^5^, Iliyan D Iliev^12^, Joonsoo Kang^9^, Jordan Voisine^2^, Josh Choi^5^, Julia Merkenschlager^13^, Jun R. Huh^5^, Katharine Block^7^, Ken Cadwell^10^, Kennidy K. Takehara^11^, Kevin Osum^7^, Laurent Brossay^14^, Laurent Gapin^15^, Liang Yang^5^, Lizzie Garcia-Rivera^1^, Marc K. Jenkins^7^, Maria Brbic^16^, Maria-Luisa Alegre^2^, Marion Pepper^8^, Mariya London^17^, Matthew Stephens^2^, Maurizio Fiusco^16^, Melanie Vacchio^3^, Michael Starnbach^5^, Michel Nussenzweig^13^, Mitch Kronenberg^18^, Myriam Croze^19^, Nalat Siwapornchai^5^, Nathan Morris^12^, Nicole E. Scharping^11^, Nika Abdollahi^19^, Nitya Mehrotra^2^, Odhran Casey^5^, Olga Barreiro del Rio^5^, Paul Thomas^20^, Peter Carbonetto^2^, Remy Bosselut^3^, Rocky Lai^9^, Sam Behar^9^, Sam Borys^14^, Sara E. Hamilton^7^, Sara Mostafavi^8^, Sara Quon^11^, Serge Candéias^21^, Shanelle Reilly^14^, Shanshan Zhang^5^, Siba Smarak Panigrahi^16^, Sofia Kossida^19^, Stefan Muljo^3^, Stefan Schattgen^20^, Stefani Spranger^22^, Steve Jameson^7^, Susan M. Kaech^1^, Takato Kusakabe^12^, Taylor Heim^22^, Tianze Wang^8^, Tomoyo Shinkawa^9^, Ulrich von Andrian^5^, Val Piekarsa^5^, Véronique Giudicelli^19^, Vijay Kuchroo^5^, Woan-Yu Lin^12^, Ziang Zhang^2^

1. NOMIS Center, Salk Institute for Biological Sciences, 2. The University of Chicago, 3. National Institutes of Health, 4. Allen Institute for Immunology, 5. Harvard Medical School, 6. Dept of Dermatology and Immunology, University of Pittsburgh, 7. University of Minnesota, 8. University of Washington, 9. UMass Chan Medical School, 10. University of Pennsylvania, 11. University of California San Diego, 12. Weill Cornell Medicine, 13. The Rockefeller University, 14. Brown University, 15. University of Colorado Anschutz Medical Campus, 16. Swiss Federal Institute of Technology, Lausanne, 17. New York University, 18. La Jolla Institute, 19. IMGT, Univ Montpellier, 20. St. Jude Children’s Research Hospital, 21. Alternative Energies and Atomic Energy Commission, Grenoble, 22. Massachusetts Institute of Technology

## MATERIALS and METHODS

### Single-cell RNAseq and TCRseq

All immgenT datasets were generated following the same SOPs (accessible at https://www.immgen.org/ImmGenT/immgenT.SOP.pdf). These datasets were obtained in the course of 66 different experiments, each experiment corresponding to one or a few encapsulation runs. Each encapsulation was numbered with an “IGTx” identifier used in this paper. This IGT code is used throughout to track individual datasets, and starts the unique cell identifier (IGT.cellID) used as the linking key between immgenT TCR or Gene Expression (GEX) data.

### Mice

As detailed and tabulated in (immgenT-Cosmo) most mice used in the immgenT project were C57BL/6 (hereafter B6; RRID:IMSR_JAX:000664) males and females originating directly from the Jackson Laboratory, with a minority of experiments performed on B6 mice maintained in colonies of immgenT participating labs (mostly for transgenic or KO experiments). Two independent “one-mouse” datasets (IGT7-9 and IGT100-101) were generated by harvesting multiple organs from a single six week-old B6 mouse. MHC-II variant mice used for IGT91/92 and IGT102/103 include the MHC-deficient Ab° line^80^ and Eα16 transgenic mice on the B6 background which express the H2-E molecule in all MHC-II+ cell-types^81^ maintained at the Jackson Laboratory. “E-only” mice (**Fig. 10**) were generated by intercrossing the Eα16 and Ab° lines. A few thymocyte samples were generated from *Tcra*-deficient Jax Strain #002116, obtained from the Jackson Laboratory. Experiments by the Core Team at Harvard Medical School were performed under following HMS IACUC protocol IS00001257.

### Sample preparation and staining

Briefly, single cell suspensions were labeled with fluorescent antibodies for cell sorting and with DNA-coded tags (TotalSeq-C) for sample identification, sorted and encapsulated on a 10X Genomics Chromium instrument. We realized that the optimal procedure to avoid contamination with hashtags and CITEseq reagents between samples was to label the cells with barcoded reagents prior to the flow cytometry step (cell washing with centrifugation was never as effective). Single-cell suspensions from tissues were obtained by mechanical dissociation, or enzymatic dissociation for some tissues, and stained with fluorescent antibodies and different TotalSeq-C Anti-Mouse Hashtags (TotalSeq-C Anti-Mouse Hashtags (Biolegend #155861-155879) in staining buffer (Phenol Red-free DMEM, 2% FCS and 10mM HEPES) for 20 min at 4°C in the dark.

For most samples, whole T cells were sorted with anti-CD3, in some cases further restricted to CD4+, CD8-, or CD8+ cells. Specific populations were sorted as described in the accompanying (**immgenT-Cosmo**) paper.

### Single-cell RNA, TCR and TotalseqC Library Preparation

Single-cell RNA sequencing was performed using the 10x Genomics 5’ v2 or v3 platform with Feature Barcoding for Cell Surface Protein and Immune Receptor Mapping, adhering to the manufacturer’s guidelines (CG000330). After cell encapsulation with the Chromium Controller, reverse transcription and PCR amplification were performed in the emulsion. From the amplified cDNA library, smaller fragments containing TotalSeq-C-derived cDNA were separated for Feature Barcode library construction, while larger fragments containing transcript-derived cDNA were preserved for TCR and Gene Expression library generation. Library sizes for both cDNA fractions were evaluated using the Agilent Bioanalyzer 2100 High Sensitivity DNA assay and quantified with a Qubit dsDNA HS Assay kit on a Qubit 4.0 Fluorometer. After enzymatic fragmentation and size selection of the cDNA, the library was ligated to an Illumina R2 sequence adapter and indexed. The three libraries were pooled based on molarity in the following proportions: 47.5% RNA, 47.5% Feature Barcode, and 5% TCR. The pooled libraries were sequenced on an Illumina NovaSeq S2 platform (100 cycles) using the 10x Genomics specifications: 26 cycles for Read 1, 10 cycles for Index 1, 10 cycles for Index 2, and 90 cycles for Read 2.

### Primary data processing

Gene and TotalSeq-C antibody (surface protein panel and hashtags) counts were obtained by aligning reads to the mm10 (GRCm38) mouse genome using the M25 (GRCm38.p6) Gen-code annotation and the DNA barcodes for the TotalSeq-C panel (see (immgenT-Cosmo) manuscript). Alignment was performed with CellRanger software (v7.1.0, 10x Genomics) with default parameters. Cells were identified and separated from droplets with high RNA and TotalSeq-C counts by determining inflection points on the total count curve, using the barcodeRanks function from the DropletUtils package.

Sample demultiplexing was performed using hashtag counts and the HTODemux function from the Seurat package (Seurat v4.1). Doublets (droplets containing two hashtags) were excluded, and cells were assigned to the hashtag with the highest signal, provided it had at least 10 counts and was more than double the signal of the second most abundant hashtag. Hashtag count data were also visualized using t-SNE to ensure clear separation of clusters corresponding to each hashtag. All single cells from the gene count matrix were uniquely matched to a single hashtag, thereby linking them unambiguously to their original sample.

Cells meeting any of the following criteria were excluded from the analysis: fewer than 500 RNA counts, dead cells with over 10% of counts mapping to mitochondrial genes, fewer than 500 TotalSeq-C counts, or positivity for two isotype controls (indicating non-specific To-talSeq-C antibody binding). Non-T cells were excluded based on the expression of the MNP gene signature, B cell signature, ILC gene signature, and the absence of T cell gene signature (score calculated using AddModuleScore_UCell)

### Gene expression data and cell annotation

The complete analysis of the gene expression data, determination of cell identities and partition into lineages (level1) and clusters (level2) is fully described in the accompanying immgenT papers (immgenT-Cosmo). Briefly, all immgenT gene expression data were integrated using the SCVI.TOTALVI model (v1.2.0)^82^ with the IGT id as a batch parameter. Dimensionality reduction was performed using the pymde.preserve_neighbors() function with default parameters [https://pymde.org/citing/index.html]. Cell clustering was carried out using the FindClusters() function in Seurat in the TOTALVI latent space.

Dimensionality reduction and 2D projection throughout the immgenT project use Minimal Distortion Embedding (MDE)^83^ rather than the more common UMAP, because MDE explicitly minimizes distance distortion and preserves relative density, thus better preserving local and global structure

Main lineage annotation (CD4, CD8, Treg, gdT, Tz, DN, DP) was done using protein and RNA expression of *Cd3e, Trbc1, Trbc2, Cd4, Cd8a, Cd8b1, Foxp3, Zbtb16, Trgc1, CD3, TCRB, THY1.2, CD4, CD8A, CD8B, CD62L, CD44, TCRGD, TCRVG1.1, TCRVG2, TCRVG3*. The initial annotation was based on the clusters. From CITEseq data, 1% were found initially mis-annotated (e.g. CD4 with CD8) and corrected. Within each lineage, we used the same initial TOTALVI latent space to recluster the cells, with further refinement or merged based on silhouette analysis, coherence in sample distribution, gene expression signatures, protein expression.

To match TCRs with the phenotype of the cells expressing them, the complete cell annotation file GSE297097_annotation_table_20250505_IGT1_104.csv.gz (accessible at GSE297097) was used, with each cell’s unique IGT.cellID identifier as a primary key. Cell coordinates on the reference immgenT MDE plots (whole data or lineage-specific) can be retrieved from the GSE297097_mde_incremental_level1_All_data.csv.gz and GSE297097_mde_incremental_level2_{$Lineage}.csv.gz files (accessible at GSE297097).

### TCRseq data pipeline

#### Contig assembly

Data from the TCR sequencing data were first processed using CellRanger v8.0.1 (10X Genomics). Raw BCL files were converted to FASTQ format using the CellRanger mkfastq command. TCR contigs for the alpha and betas chains were assembled using the CellRanger VDJ pipeline (run with default parameters) with the C57BL/6 mouse reference genome from IMGT (Release 202416-4). CellRanger, in versions v7.2.0 and above, selects the in-frame contig with the greatest number of reads as the primary “alpha” and beta”. In many (but not all) cells a second “alpha2” and “beta2” contig that represents a second in-frame transcript or an unproductive join, is generated and was considered here. In rare cells, more contigs are generated, usually at low read numbers, possibly representing excision circles as proposed^65^.

These were not considered and are not included in the SummaryTable. Contig FASTA files were all keyed to the immgenT-wide IGT.cellID identifiers (the IGT number followed by the 16-base barcode of each cell), and are available (under accession number GSE297097).

In some experiments, to cross-check the accuracy of the TCR contig assemblies generated by CellRanger, we independently generated TCR contigs from the raw data using MiXCR (v.4.7.0) ^52^, using the same IMGT mouse reference genome library (C57BL/6 background) for contig reconstruction.

#### TCR sequence parsing and identification of V, D and J elements

FASTA files of the assembled contigs were analyzed using IMGT/HighV-QUEST (version 1.9.5, hereafter V-QUEST^84^ with the *Mus musculus* (C57BL6/J) reference directory (release 202416-4), using default parameters, plus the “Search for insertions and deletions in V-REGION”. A SummaryTable was generated from the V-QUEST output folders using a custom Jupyter script (available on GitHub at https://github.com/immgen/TCR) to extract the sequence functionality, V and J composition, junction sequences (AA and nucleotides), and predicted N-region nucleotides for all chains (alpha, beta, alpha2, beta2) in each cell. Additionally, the SummaryTable was enriched with metadata including organ source, genetic variation, cell type annotations, cluster assignments, coordinates for the Seurat UMAP generated for each IGT, and coordinates for the immgenT-wide integrated MDE (see immgenT-Cosmo).

#### “Simplification” of ambiguous V gene name assignments

For a sizeable number of cells, there was ambiguity in calls of the exact *TRAV* gene used by a cell, owing to duplications and triplications of some genes of the *TRAV* locus in B6 mice. In several analyses, especially those pertaining to cellular selection of the repertoire or to association with the TCRβ chain, these ambiguities were not relevant, but they needlessly complicated the representations and power. In such cases, we thus adopted a simplified assignment of *TRAV* genes (for instance, *TRAV3N-3* and *TRAV3D-3* were both assigned to *TRAV3-3* in the “simplified” tables used). These edits are listed in **Table S12**. Some *TRBJ* assignments by V-QUEST were ambiguous, and were resolved as: 68 cases of “*TRBJ1-1* | *TRBJ1-2*” assigned as “*TRBJ1-1*”; 13 instances of “*TRBJ2-4* | *TRBJ2-5*” assigned as “*TRBJ2-5*”

#### Determination of V and J genes usage frequencies

Tables of *V* or *J* gene usage were generated from the SummaryTable after indexing according to cell-type (retrieved from GSE297097), organ, experiment or sample, and displayed as comparative scatter plots or heatmaps generated in Morpheus (Broad Institute; RRID:SCR_017386). For plots of VaJa pair usage ordered according to genomic position based on gene positions of *Mus musculus* C57B6L/6J in IMGT repertoire (https://www.imgt.org/IMGTrepertoire/LocusGenes/index.php?species=house_mouse#GenePositions-tab-pane), heatmaps were generated in Jupyter Notebook using Seaborn package (v.0.13.2; DOI:10.21105/joss.03021).

#### Public specificities

To identify public specificities, datasets involving TCR transgenic mice were excluded. We first defined TCRαβ clonotypes as sharing of both chains at the amino-acid level, as a character string combining alpha.vgene, alpha.junction, beta.vgene and beta.junction (we found that adding the J segments did not further split these clonotypes, as the J identity is represented in the junction sequence). Only instances involving sharing by cells from different IGT datasets were considered further, to avoid simple cases of clonal expansion - this strict criterion may under-represent public specificities, as many IGT datasets involve several independent mice. These instances of shared clonotype at the protein level were then further split into those that only shared protein identity vs the smaller group that also shared full identity at the nucleotide sequence.

We suspect that some contamination may have occurred accidentally during pooling and sequencing of the TCR libraries, from IGT3 into the IGT44 library, and from IGT100/101 into IGT91/92. These contaminations appeared as cells from different datasets that shared a TCRαβ clonotype and also shared the same 10X cell barcode across experiments, a vanishingly improbable occurrence. We thus systematically excluded TCR clonotypes shared between these datasets as supporting a public specificity.

#### Analysis of non-canonical joins

To identify extreme junctions’ length of rearranged productive *TRA* and *TRB*, only sequences from non-transgenic mice, and with a correctly identified *C* gene were included (the *C* gene was identified by detecting the inner primer sequence in exon 1 of the respective *C* gene). We employed an empirical probability approach based on the distribution of observed values of junction lengths computed by V-QUEST. Extreme junctions’ length was defined using percentile-based thresholds of 0.5^th^ and 99.5^th^ percentile for the lower and upper thresholds respectively. Rearranged productive TRA and TRB sequences with unusually short (<10 AA) or long (>18 AA) junctions were then extracted for detailed analysis. Sequences from the same experiment sharing identical *V* and *J* genes and nucleotide junctions were collapsed. To test whether extreme junction lengths in one chain were associated with correspondingly shorter or longer junctions in the paired chain we evaluated their association using a Chi-square test. In this set of sequences, *V* and *J* gene assignments were manually verified, along with the number of nucleotides trimmed at the 5’ end of the *V* region, both ends of the *D* region when identifiable and 3’ end of the *J* region, and the addition of N and P nucleotides.

To investigate unusual rearrangements, all sequences were analyzed using BLASTn (v2.16.0) against the *Mus musculus* (taxid:10090) core nucleotide database (core_nt). Sequences aligning to two *V* or *J* genes, even partially, at distinct positions, were manually reviewed to confirm gene assignments and examine the intervening sequence and putative mechanisms (e.g., presence of a putative RSS heptamer). Trans-rearrangements between *TRA* and *TRB* genes were identified by blasting *TRBV/TRBJ* against *TRA* sequences and *TRAV/TRAJ* against *TRB* sequences, respectively, and *TRAC* or *TRBC* genes were confirmed via BLAST.

The N region of all sequences was examined to detect stretches of ≥5 nucleotides matching *TRDD* genes in *TRA* sequences involving *TRAV* genes. For rearrangements involving *TRDJ1*, BLASTn was used to verify whether the associated *C* gene belonged to *TRA* or *TRD*. Similarly, sequences were checked for the presence of the second *TRBD* D-REGION within the N region. Finally, unusually long N (≥13 nt) nucleotide stretches in which no *TRBD* or *TRDD* gene could be identified were isolated and blasted against the reference *Mus musculus* (taxid:10090) core nucleotide database (core_nt) to detect their potential genomic origin.

#### Refinement and analysis of N region diversity

To consider potential sequencing errors and/or potential somatic mutations, the N region was corrected when it contained ≥ five consecutive nucleotides matching the IMGT *J* gene reference before a mismatch, allowing up to two mutations in the 5’ *J* in V-QUEST analysis. Annotation in the SummaryTable also accounted for potential artifacts and ambiguous assignments. Potential trans-rearrangements between *TRA* and *TRB* were flagged as “Rearrangement TRAV with TRBJ|TRBC” or “Rearrangement TRBV with TRAJ|TRAC”. If the N-REGION assignment was uncertain due to unusual rearrangement, its sequence was replaced with the label “No call freaks”. For alpha chains, N-REGION with potential *TRDD1* or *TRDD2* elements were labeled “| Possible delta D”. The | symbol was used to separate the N nucleotides located before and/or after the potential *TRDD* segments. For beta chains where the D gene could not be identified because of the short number of nucleotides (less than three), N-REGION were labeled “| Possible D” in the .vd.nregion column, as a few N nucleotides might come from a *D* gene.

Sequences missing the conserved Cys-104 or Phe/Trp-118 residues were further analyzed using BLASTn (v2.16.0). Alignments were compared with full-length *V* and *J* genes to detect over-trimming, and sequences were annotated as “No call overtrimmed” in the SummaryTable. The preference for addition of individual bases among N nucleotides was calculated from single-base additions, and was used in simulations of random generation of N.nucleotide x-mers for **Fig. 4F**.

#### TCR Generation Probability Analysis in OLGA

To quantify the statistical rarity and public sharing potential of T cell receptors, we computed the generation probability (Pgen) for each unique CDR3 sequence using OLGA (v1.2.4)^29^. The input data consisted of amino-acid and nucleotide junction sequences with their corresponding *V* and *J* gene assignments, extracted from the full SummaryTable. Pgen values were calculated using the standard mouse generative model, which employs a dynamic programming algorithm to sum over all possible V(D)J recombination events, including insertions and deletions, that could produce a given sequence. These values were then converted to Pgen scores through negative log transformation (-log10) to facilitate display.

#### Alternative splicing

To explore the presence of trans-splicing in the beta chain between the L-PART1 of a gene and the V-EXON of the downstream Variable gene, all sequences assigned to each identified *V* gene were extracted from the input FASTA files and analyzed using BLASTn (v2.16.0). For each gene, sequences were blasted against the alternative splicing sequence of its V-EXON and the L-PART1 of the upstream gene. Sequences were considered true alternative splicing events if they aligned with both the V-EXON of the gene and the full length of the L-PART1 (100% identity) of the upstream gene. In the case of the genes in the *TRBV13* and *TRBV12* family, we blasted the L-PART1 of one gene against V-EXON from all downstream genes from these two families. Regarding the *TRBV13-2* gene, the SummaryTable columns ‘beta.vgene’ and ‘beta2.vgene’ were annotated as follows: ‘true’ when L-PART1 is from *TRBV13-2*, ‘chimeric (chim)’ when the L-PART1 is from *TRBV12-2*, and ‘uncl’ for partial alignment of the L-PART1 from *TRBV12-2* (**Table S4A).**

#### Search for somatic mutation

We focused the search of potential somatic mutations exclusively on sequences from pure JAX C57BL6/J mice. We first extracted sequences with mismatches in the V-REGION, J-REGION, and C-REGION by comparing them to the IMGT reference directory. This extraction was based on output files from V-QUEST for the V-REGION and J-REGION, and on BLAST analysis for the C-REGION. A set of stringent filtering criteria was applied on these sequences to exclude mismatches that might be sequencing errors and not somatic mutations. We analyzed only productive sequences with identified primer sequences in the C region as well as sequences covering the full V-DOMAIN (V-D-J-REGION or V-J-REGION). When identified, chimeric sequences were also excluded from the analysis, as well as sequences with the genes *TRAV9N-4, TRAV9-1, TRAV9D-1,* and *TRAV13D3*, since mismatches could result from the rearrangement of pseudogenes *TRAV9D-4, TRAV9N-1*, and *TRAV13-3,* which are not included in the V-QUEST reference set. For each mismatch, at least five reads were required to cover the position and had to be beyond the first 10 nucleotides on 5’ of the contig. The mean quality score of the reference reads has to be superior to 20 when reference reads cover the mismatched position, and the mean quality score of the reads containing mismatches has to be above 30. Finally, mismatches located at the 3’ end of the V-REGION and 5’ end of the J-REGION that are due to trimming are also not retained. Additionally, for the V-REGION, we extracted the number of UMIs per mismatch and considered only those positions with more than one UMI for the mismatch as true somatic mutations, as sequencing errors are less likely to occur at the same position in multiple independent molecules.

Mutations at the same position in the same genes and with the same nucleotide mutated in one sample were collapsed and the mutation rate (per 100bp) was calculated for each region: V-REGION, J-REGION, and C-REGION of TR genes. Mutation rates were also calculated for some other T receptor genes (e.g. *Cd4, Cd8a, Cd28 and Cd3e*) based on the scRNAseq data from the secondary lymphoid organs, peritoneal cavity, and blood T cells (IGT10, IGT11 and IGT12). The mutations were counted when the base quality was over 30, and the position was covered by at least five reads. Mutations found in only one UMI in a cell were ignored as likely reflecting a PCR error.

#### Recurring rearrangements at VaJa, AJ joins

To eschew distortions from clonal expansion, we used as input combined immgenT data in which all clonal expansions within a given dataset (identified as full sharing at nucleotide level on both chains) were collapsed down to a single instance (hereafter “datc”, n=309,806 cells). Within data, we calculated for each VaJa pair its total occurrence (“count”), and the frequency of the most common amino-acid junction sequence (“main3p”). Friedman’s SuperSmoother an adaptive smoothing algorithm that applies running linear smoothers at multiple spans, was used to plot estimate the main relationship between count and main3p value for all VaJa pairs in **Fig. 5A**. To estimate the significance of the AJ joints discovered, which are essentially outliers in the count vs main3p distribution, we computed the probability of the main3p value of each VaJa relative to the mean and standard deviation of main3p for other VaJa joins that fall in the same scanning window of count values (pnorm() function).

#### Analysis of nonproductive joins

Here, all non-transgenic cells were used. The CellRanger algorithm typically orders the most abundant productive allele in the first contig, and places in the second contig (alpha2 and beta2 in the SummaryTable) the unproductive allele (or a second productive allele). Unproductive joins on alpha2 or beta2 chains were flagged if “functionality” was called NOT “productive” by V-QUEST, AND a *TRAV* (or *TRBV*) was positively identified. Matrices were generated by simple counting of unproductive/nonproductive Vbi-Vbj pairs, and normalized for **Fig. 6A** by dividing by the column-mean (=average usage of each *TRBV* in productive joins) before log2 transformation and display in Morpheus. The same calculation was performed for *TRAV* productive/unproductive pairs (**Fig. 6B**)

#### Allelic exclusion

Input data left out datasets from thymus or TCR transgenic mice. All four chains contigs output by the CellRanger process were considered (alpha, alpha2, beta and beta2 in the SummaryTable) and a cell was categorized as ab, aab, abb or aabb if these chains were classified as “productive” by V-QUEST and had a positively identified *TRAV* (or *TRBV*).

#### TCR meta-clonotype analysis in CoNGa

Integrated analysis of single-cell transcriptomes and TCR was performed using the CoNGA (Clonotypes Neighbor Graph Analysis) framework (version 0.1.2)^32^. The input data consisted of matched scRNA-seq gene expression matrices and clonotype data based on the SummaryTable, including IGT.cellID, amino-acid and nucleotide CDR3 sequences with their corresponding *V* and *J* gene assignments, and the numbers of reads and UMIs. CoNGA identifies correlations between gene expression (GEX) profiles and TCR sequence features by constructing two independent k-nearest neighbor (k-NN) graphs: one based on principal components of the GEX data and the other based on TCR sequence similarity using the TCRdist metric. “CoNGA clusters” or “clumps” (statistically significant overlaps between the two modalities) were identified by assessing whether clonotypes clustered in TCR sequences also occupied similar neighborhoods in GEX profiles. We utilized the default neighborhood size (k=20) and calculated CoNGA scores to identify CoNGA clusters where GEX and TCR similarities converged significantly (p < 0.05 after Benjamini-Hochberg correction).

#### Parse Bioscience TCR dataset

To compare our results with a public dataset, we extracted TCR sequencing data from the ‘TCR Sequencing of 1 Million Mouse T Cells in a Single Experiment’ made available by Parse Bioscience (https://www.parsebiosciences.com/datasets/tcr-sequencing-of-1-million-mouse-t-cells-in-a-single-experiment/#download). Sequences were extracted from their file *‘*AIRR TSV file, and analyzed using IMGT/HighV-QUEST as above. As for the ImmgenT dataset, we generated a SummaryTable including both productive and unproductive sequences (available upon request). Cells with more than two alpha or beta chains were excluded. To reduce ambiguity in gene and allele identification, we retained only sequences with a complete J-REGION and a V-REGION and with fewer than 200 nucleotides missing from the 5’ end of the V region. This yielded a final dataset of 516,345 cells, including 123,924 with both TCRα and TCRβ.

#### Human TCR dataset

To ask if AJ TCRα are also found in human T cells, we used a dataset from human PBMC, 55,277 cells from 33 healthy control donors (median 987 cells each), sourced from a commercial vendor (Immunospot). These donors included 19 male, 14 female ranging in age from 18-64 years old. These data are listed in Table S13. For each TRAV/TRAJ pair, counts and frequencies were computed as for mice, after removing clonal expansions.

### Accompanying Web resources

#### TCR Browser (https://rstats.immgen.org/tcrbrowser/)

The immgenT Project provides for the community a web-based complement for searchable and interactive access to immgenT αβTCR data. It integrates TCR sequencing results across all immgenT experiments, processed through a standardized pipeline utilizing immunarch, tcrdist3, CoNGA, and custom analytical scripts. The portal’s capabilities are slated to expand over time, but as of this writing include two interfaces:

1. View immgenT Data tab: Access to and export of data SummaryTable for every experiment, grouped by IGT number. The main IGT table is searchable by organ, challenge, cell-type, etc. The chosen data can then be retrieved as:

a. Download a SummaryTable with per-cell TCR composition, enriched with multi-layered metadata, including: IGT and sample, organ, genetic variation and immgenT level1-lineage and level2-cluster annotations, cell-level UMAP/MDE coordinates derived from the corresponding gene expression data.
b. an html file that groups the results of TCR analyses for the chosen IGT: diversity (clonotype pie-chart, duplication plot, standard metrics computed by immunarch), *TRAV* and *TRBV* usage plots, position of the most expanded clonotypes on CD4vsCD8 and UMAP, TCRdist plots and an integrated CoNGA analysis.
2. Find My TCR tab: flexible query interface allows users to search for specific TCRs by any combination of V and J element and junction sequences (amino-acid or nucleotide) to retrieve all the TCRs that match a structure/sequence of interest.

## EXTENDED DATA FIGURE LEGENDS

**Fig. S1:**
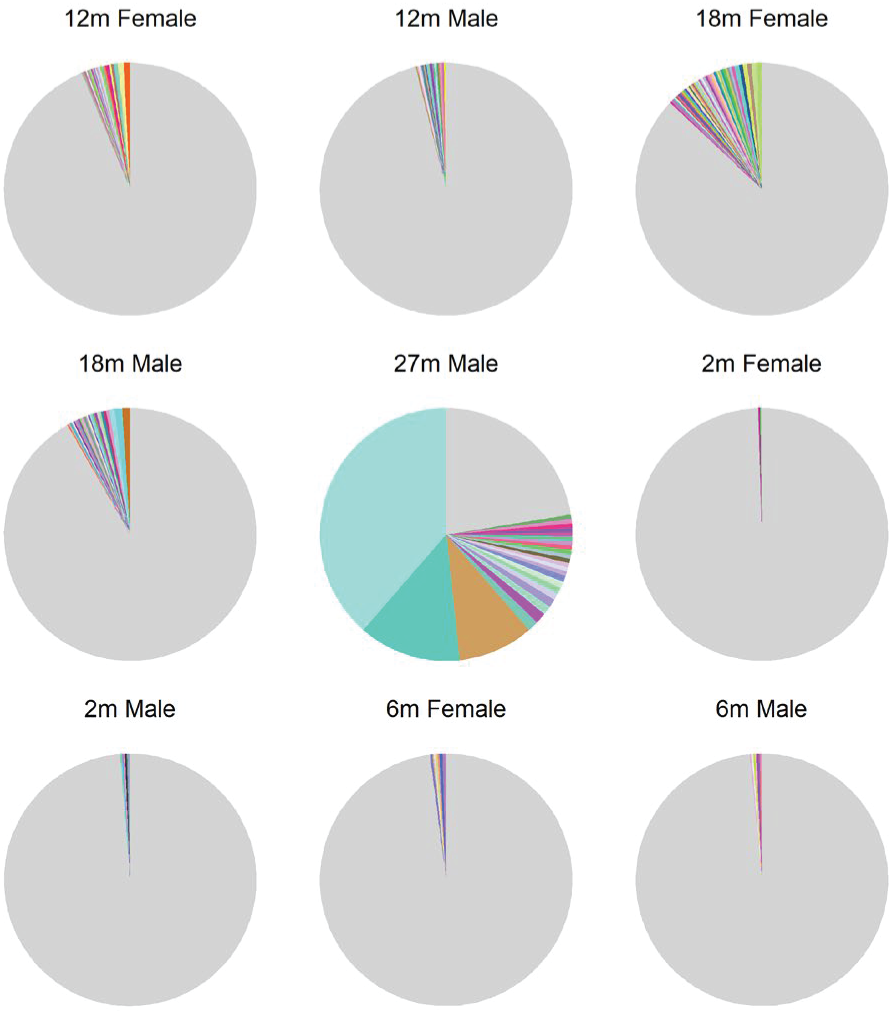
Repertoire size analysis. Pie chart of TCR clonotype distribution in adult mice of different ages (all downsampled to n-200 cells).

**Fig. S2:**
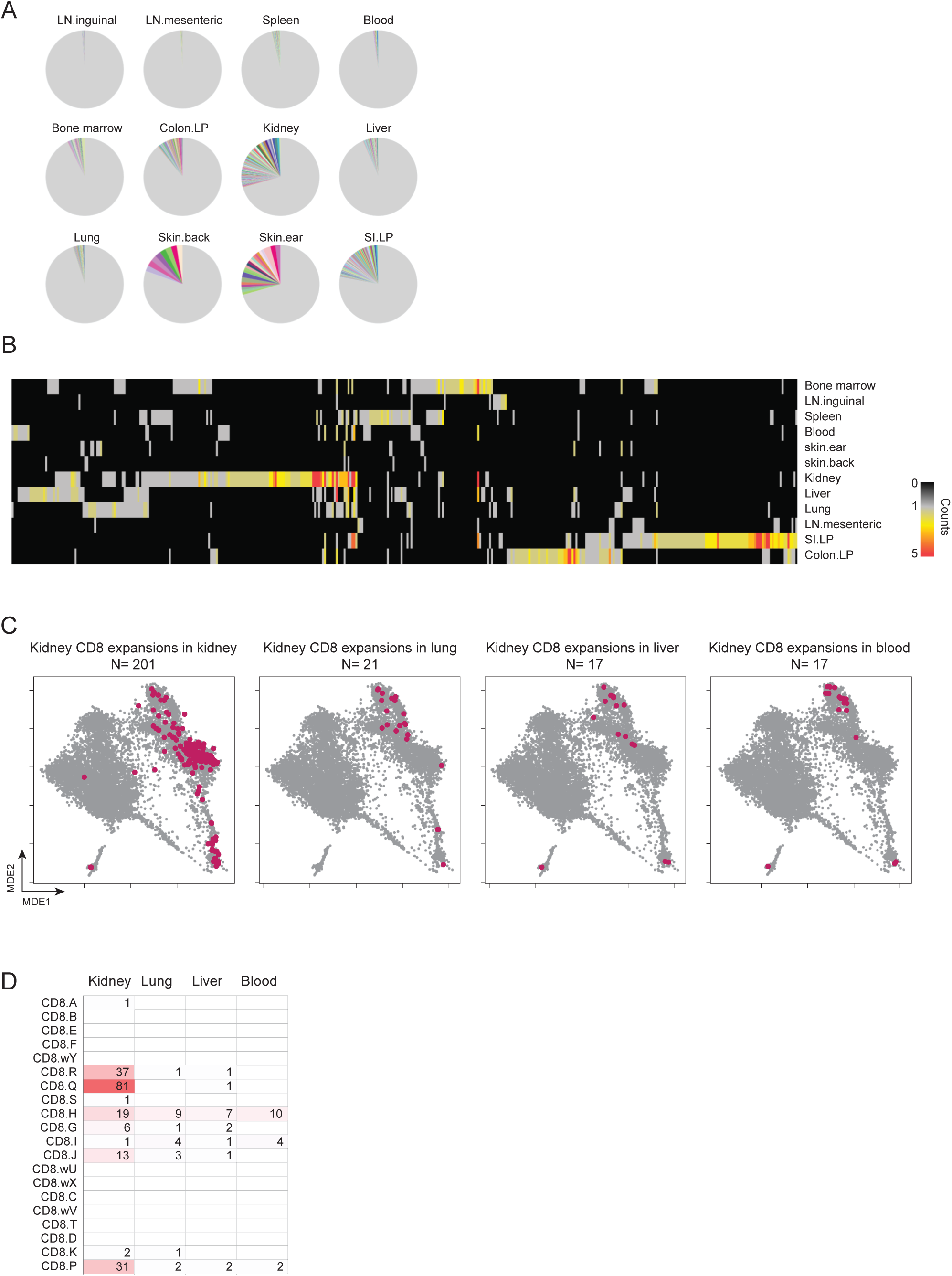
TCR repertoire distribution in one mouse. **A.** Replicate “single-mouse” experiment (IGT100/101). Pie chart of TCR clonotype distribution for different organs (all downsampled to n=100 cells). **B.** Same data as in **A**, Distribution of TCR clones detected at least twice in the organs of a single mouse, ordered by hierarchical clustering, with the cell-types shown below. On the right is indicated the total number of cells with full TCR data in each organ. **C.** Phenotype of the cells expressing the kidney-preferential clonotypes is indicated by projecting them onto MDE plot for CD8 T cells in different organs. **D.** Number of cells expressing the kidney-preferential clonotypes in different CD8 clusters, for the different organs.

**Fig. S3:**
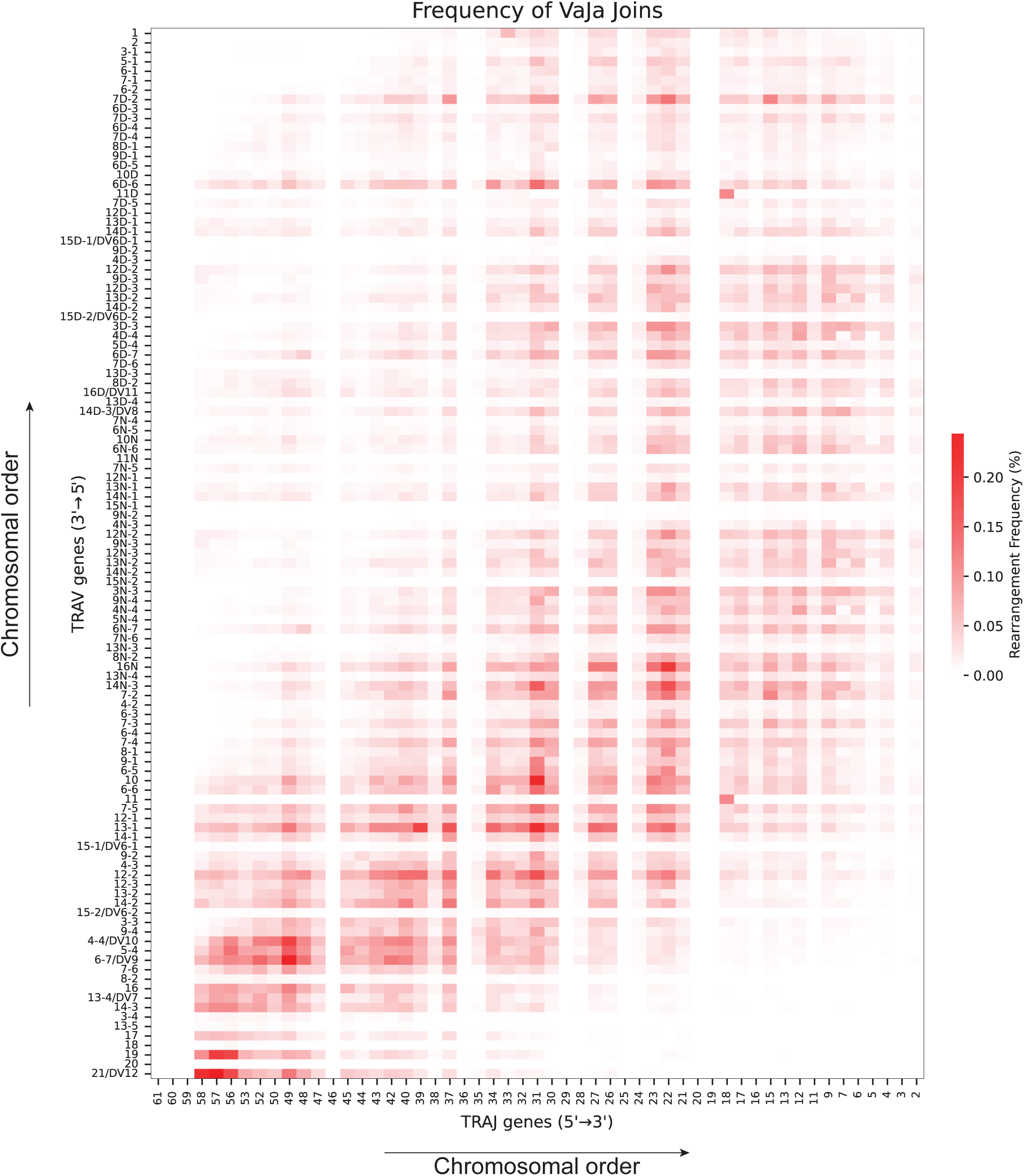
Frequency of different *TRAV/TRAJ* (VaJa) combinations. Heatmap representation, formatted as^36^, of frequency of rearrangements between *TRAJ* (x-axis) and *TRAV* (y-axis) genes, ordered according to their chromosomal position. For readibility, the “TRAV” and “TRAJ” prefixes have been omitted from gene names in the representation.

**Fig. S4:**
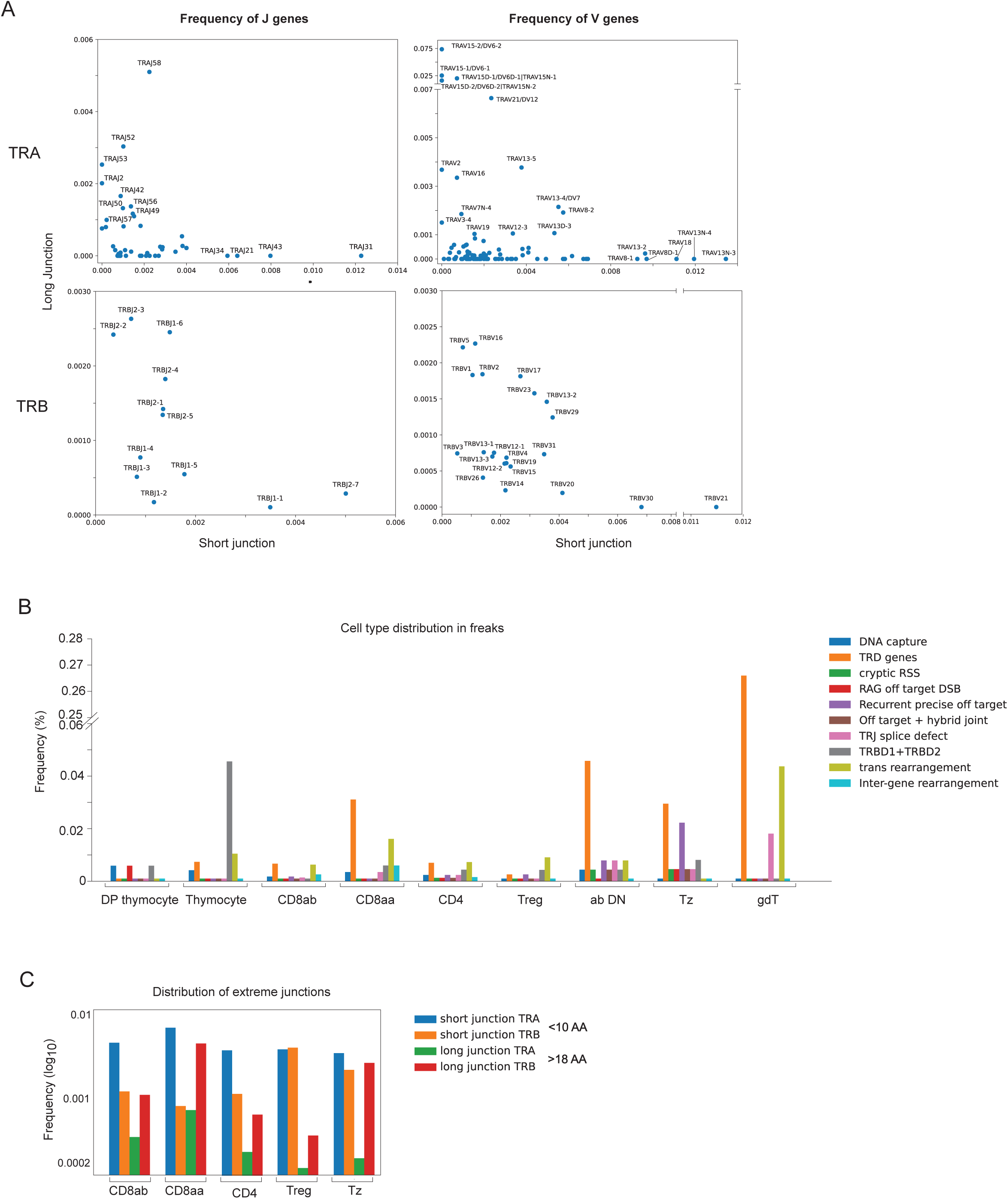
Extremely short and long junctions. **A.** Representation of *V* and *J* genes usage frequency (%) in very short (< 10 AA) versus very long (> 18 AA) TCRα and TCRβ junctions. **B.** Cell-type distribution for all types of unusual TCR VDJ recombination structures (per Fig. 4C). **C.** Distribution of extreme junctions (short <10 AA and long >18 AA) in different cell-types

**Fig. S5:**
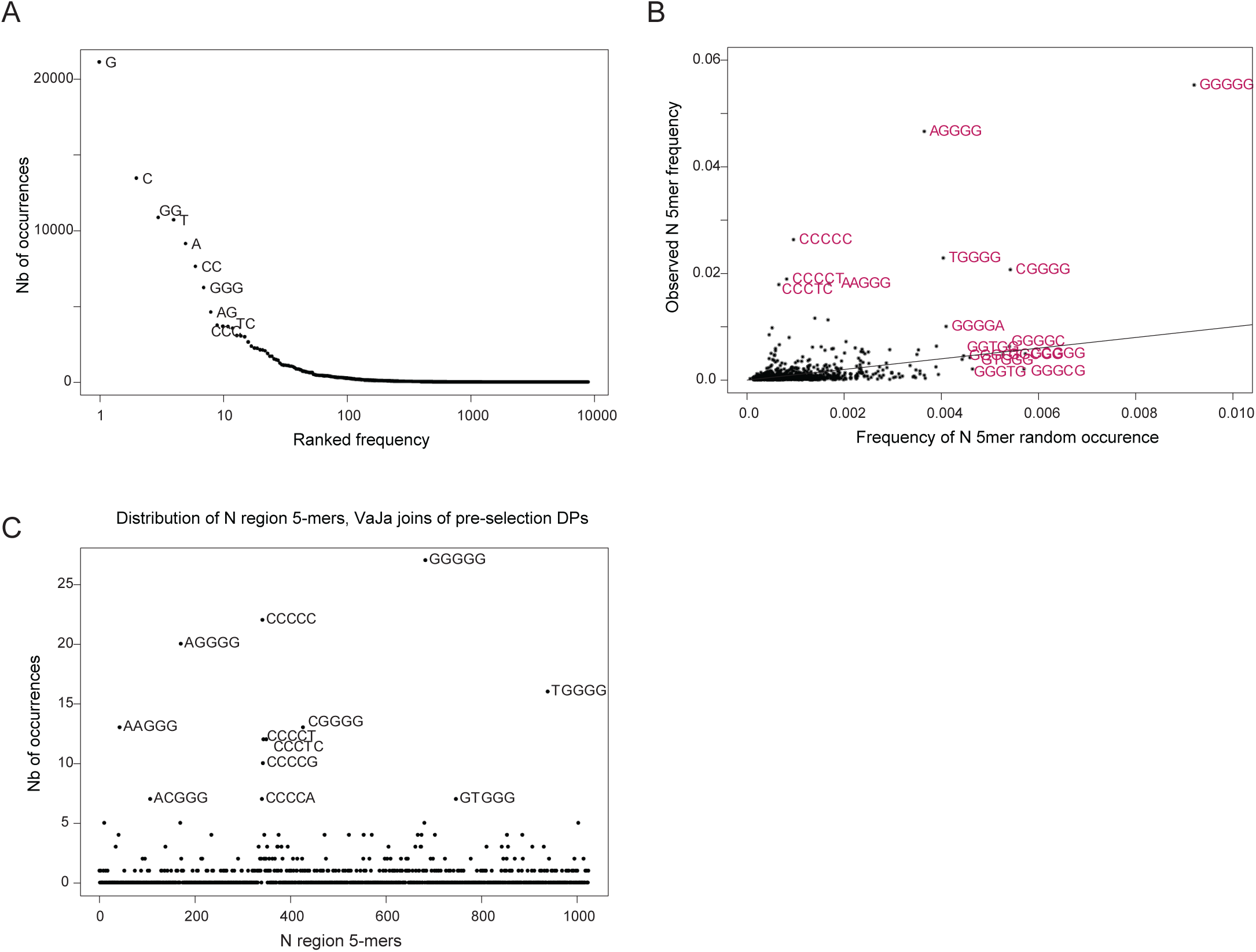
Analysis of the N-nucleotide additions. **A.** Ranked frequency of N sequences versus the number of occurrences (actual bases only shown for common mono- and di-nucleotides). **B.** Expected frequency of N region 5mers (calculated by random sampling weighted according to single-base additions) versus observed frequencies **C.** Distribution of N region 5mers in VaJa joins from pre-selection CD4+CD8+ DPs in the thymus.

**Fig. S6:**
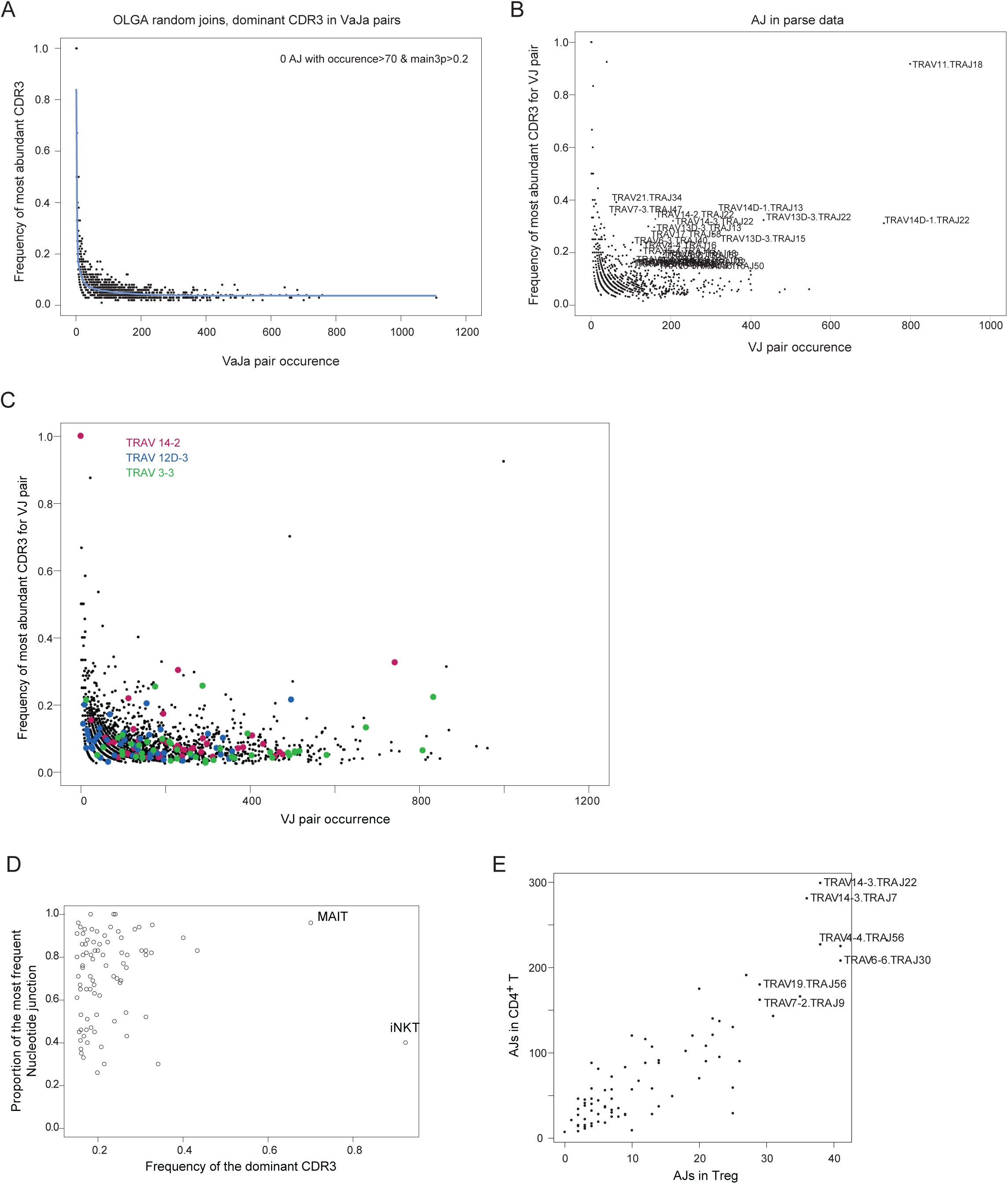
VaJa “Anomalous Joins”. **A.** For each VaJa pair generated probabilistically by OLGA (200,000 random sequences), total occurrence is plotted against the frequency of the most abundant CDR3 encoded by that pair **B.** As for A, but derived from VaJa pairs in the Parse Biosciences spleen T cells dataset **C.** For each VaJa pair, the total occurrence in the immgenT non-redundant dataset is plotted against the frequency of the most abundant CDR3 encoded by that pair. VaJa pairs that include *TRAV14-2, TRAV12D-3* and *TRAV3-3* are shown as pink, blue and green colored dots, respectively. **D.** Frequency of the dominant CDR3 for each AJ VaJa join (x-axis) is plotted versus the frequency of the most abundant nucleotide sequence for that junction. MAIT and iNKT are indicated on the plot for reference. **E.** Distribution of the number of AJs VaJa joins in Treg versus the number of AJs in CD4+ T cells.

**Fig. S7:**
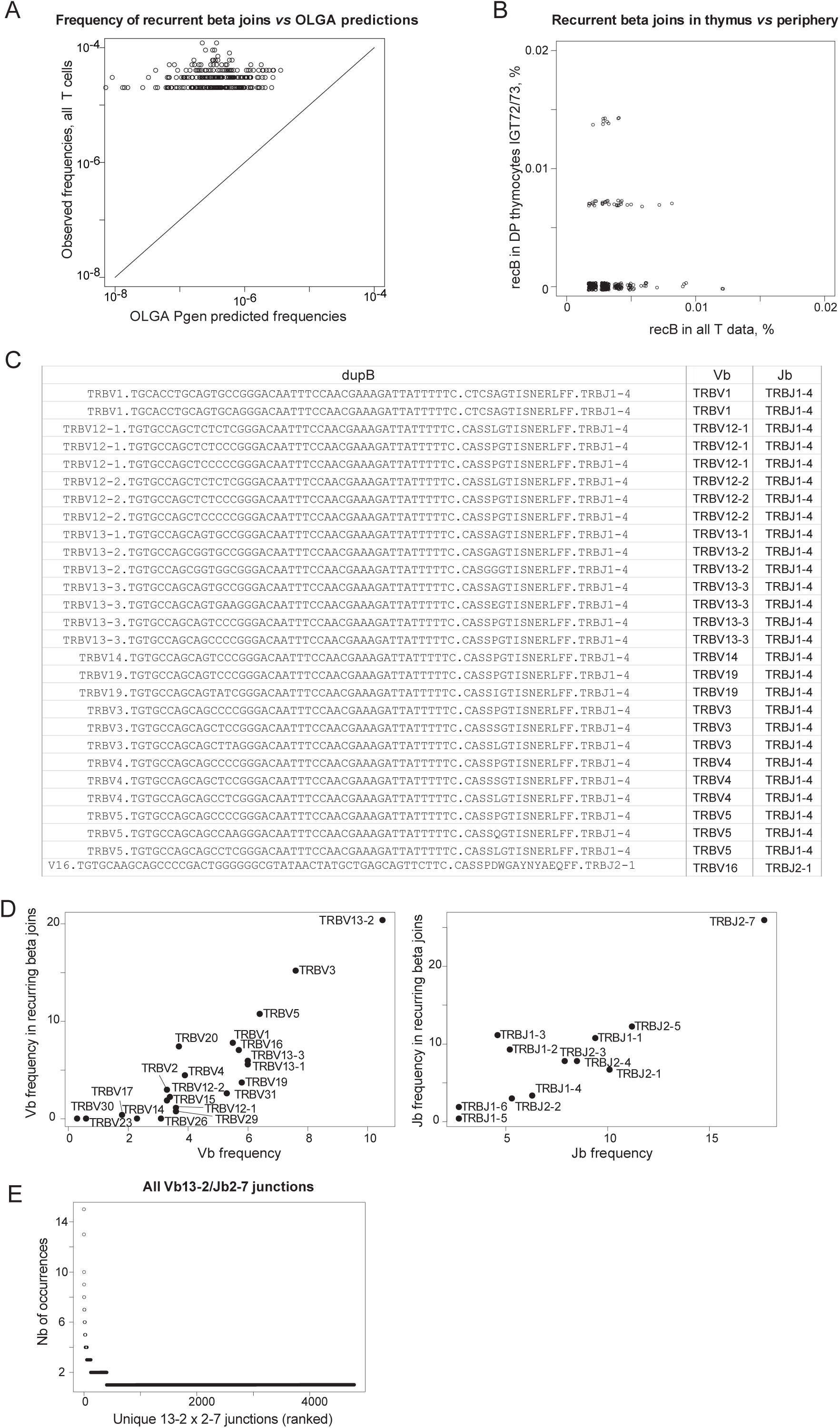
Recurring rearrangements in single-chain clonotypes. A. Comparative frequencies of the recurring TCRβ rearrangements (y-axis) plotted vs the OLGA-computed predictions of probability of generation (pGen, x-axis). B. Recurrent beta joins frequency in all T cells (x-axis) versus the frequency of beta joins in pre-selection thymic DP (y-axis). C. List of bN-clonotypes unique to CD8aa T cells. In the first column, the clonotype is described with the *TRBV* gene, the junction in nucleotides and amino-acids and the *TRBJ* gene. D. Frequency of *TRBV* genes (left) and *TRBJ* genes (right) in all data (x-axis) versus their frequency in recurring bN-clonotypes (y-axis), excluding CD8aa-specific clonotypes. E. Junction sequences involving the *TRBV13-2/TRBJ2-7* combination, ranked by number of occurrences across the non-redundant dataset.

**Fig. S8:**
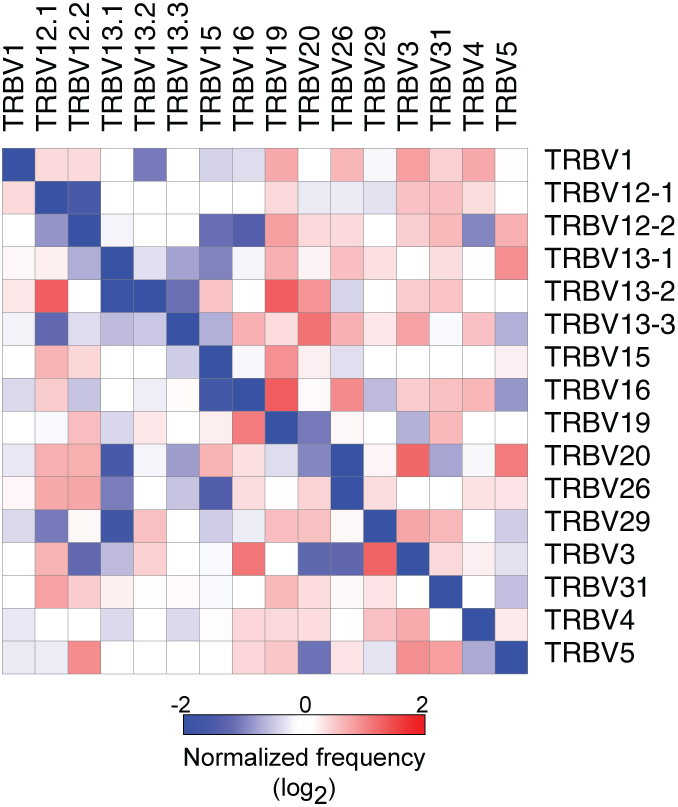
Interference between unproductive and productive joins. **A.** TCR contigs were generated independently from raw data (IGT11,12,13,14,19) using MIXCR software, and their V, D and J elements parsed and tabulated as usual with V-QUEST. The relative proportions of productive/unproductive pairs of rearranged *TRBV* in the primary and secondary alleles of each cell was counted as for Fig. 6, in cells where the contigs yielded the same assignment for contigs derived from CellRanger and MIXCR. Values are normalized vs the mean usage of each *TRBV*.

**Fig. S9:**
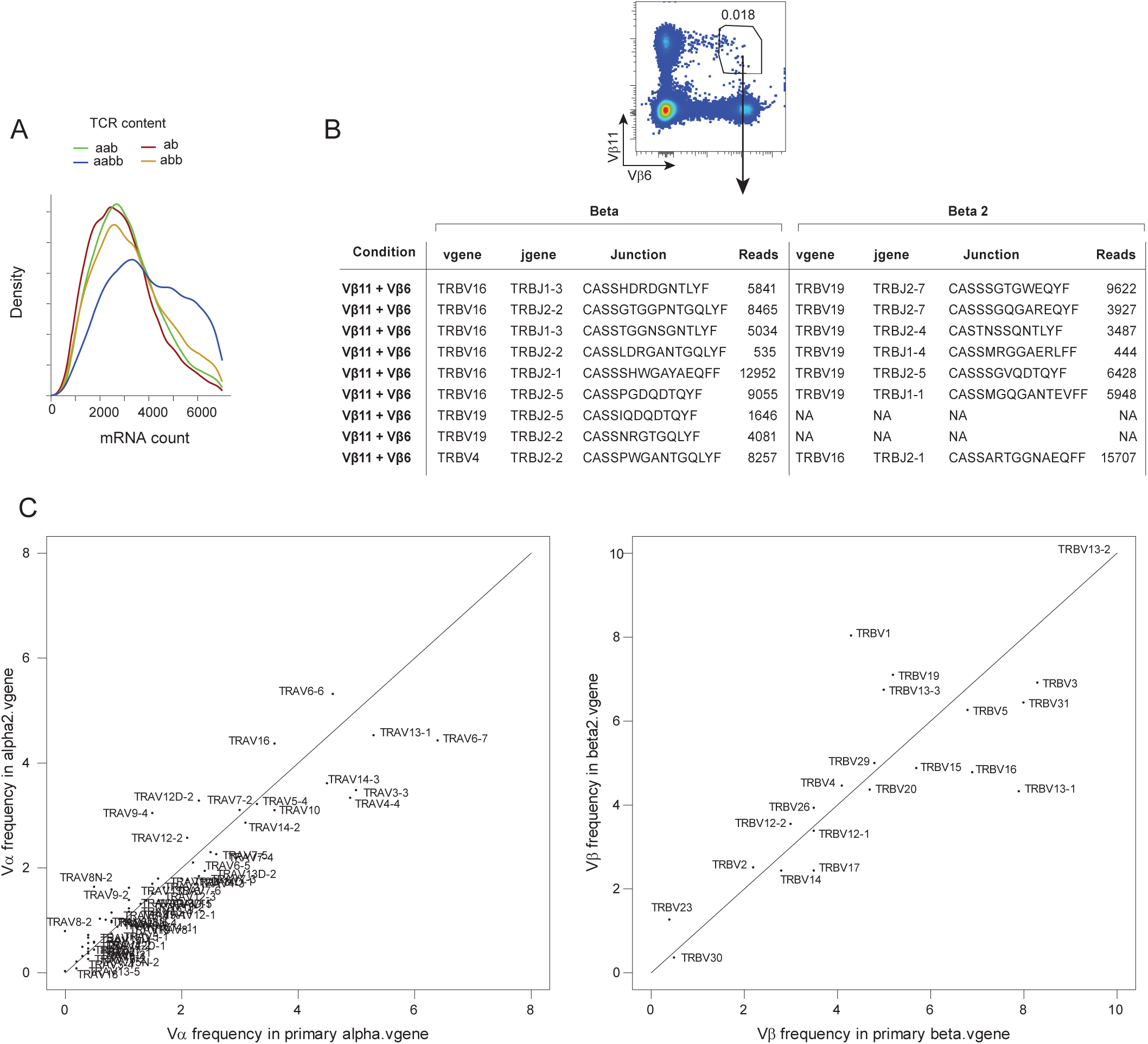
Allelic exclusion for TCRα and TCRβ. **A.** Distribution of sequencing read depth for TCR in cells with dual-α, dual-β, dual-αβ or single-αβ cells. **B.** Validation of dual-β cells: splenic T cells were sorted as dually staining with anti-Vβ6 and anti-Vβ11 mAbs, and their TCRβ composition determined in a TCRseq run. **C.** Overall distribution of *TRAV* (left) and *TRBV* (right) gene frequency (%) in the primary productive join (x-axis) versus the second productive join (y-axis) of dual-TCR cells.

**Fig. S10:**
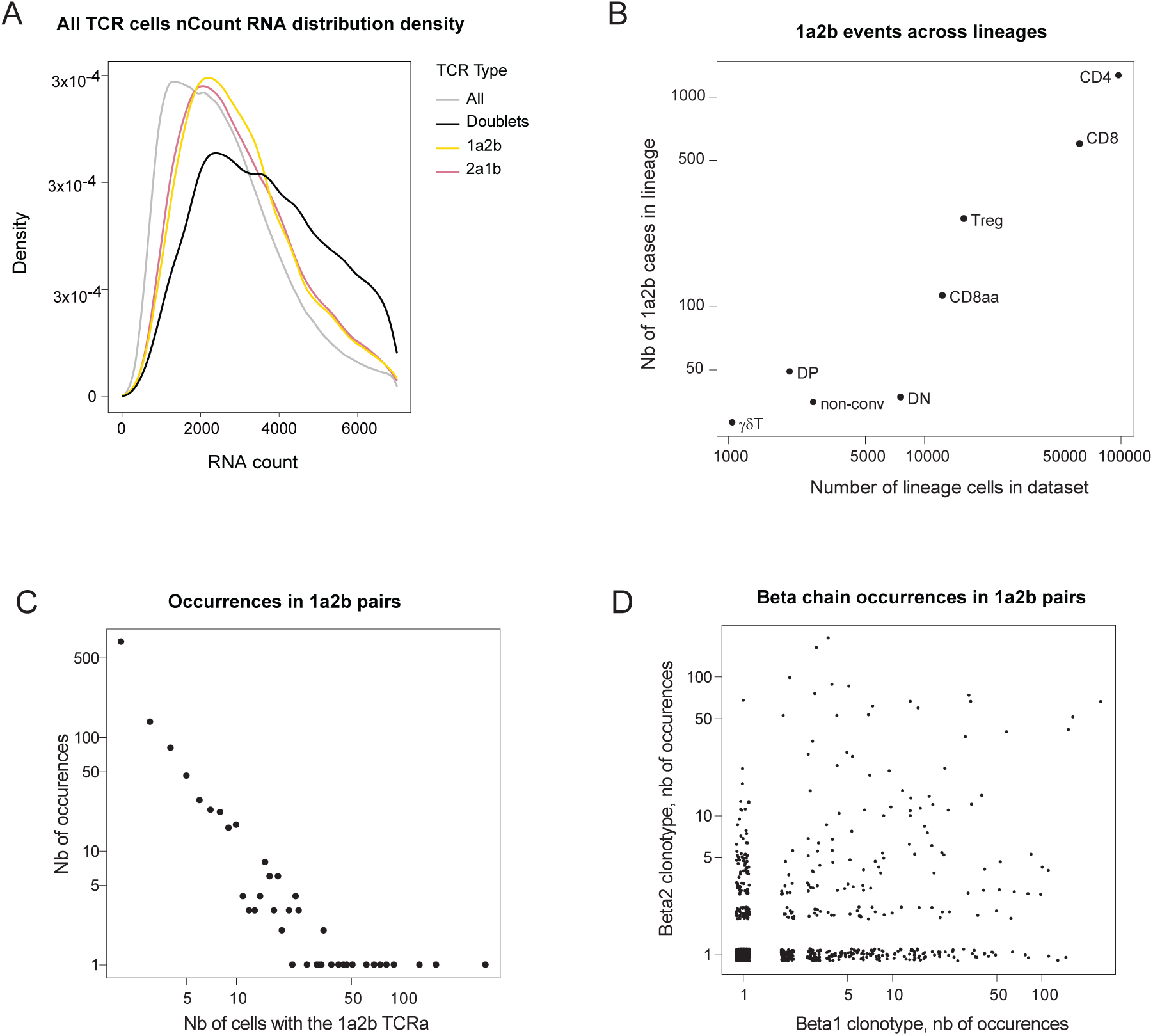
“1a2b”: same TCRα paired with different TCRβ clonotypes. **A.** Distribution of sequencing reads for TCR in all cells, in cells called as doublets by the processing pipeline, or in cells which share the same TCRα clonotype but have two different TCRβ chains (1a2b events), or cells which share the same TCRβ clonotype but have two different TCRα chains (2a1b) **B.** Distribution of 1a2b cells in different lineages (note that these pairs of cells mostly belong to the same lineage). **C.** Frequency distribution of the TCRα clonotypes that belong to 1a2b events. If the repeated presence of 1a2b TCRα clonotypes were due to contamination by ambient RNA, they would be found frequently in the sample. This may be the case for a small minority of events (78/1132 a1b2 events with TCRα clonotype incidence >20) but most (830 instances) are detected only two or three times. **D.** Number of cells that express the beta1 or the beta2 clonotypes associated with each 1a2b event. If the 1a2b events were due to contamination by TCRβ clonotypes in ambient RNA, this would be manifest as high frequency of one or the other TCRβ clonotype. This may be the case for a minority of events (representation of beta1 or beta2 clonotypes >10 times, with other occurring only once, in 66/1132 1a2b events) but not for the majority of 1a2b (752/1132 1a2b cases have the two beta clonotypes represented in only one or two cells). Note also that 92/1132 1a2b cases show clonal expansion of both TCRβ clonotypes together with the same TCRα.

**Fig. S11.**
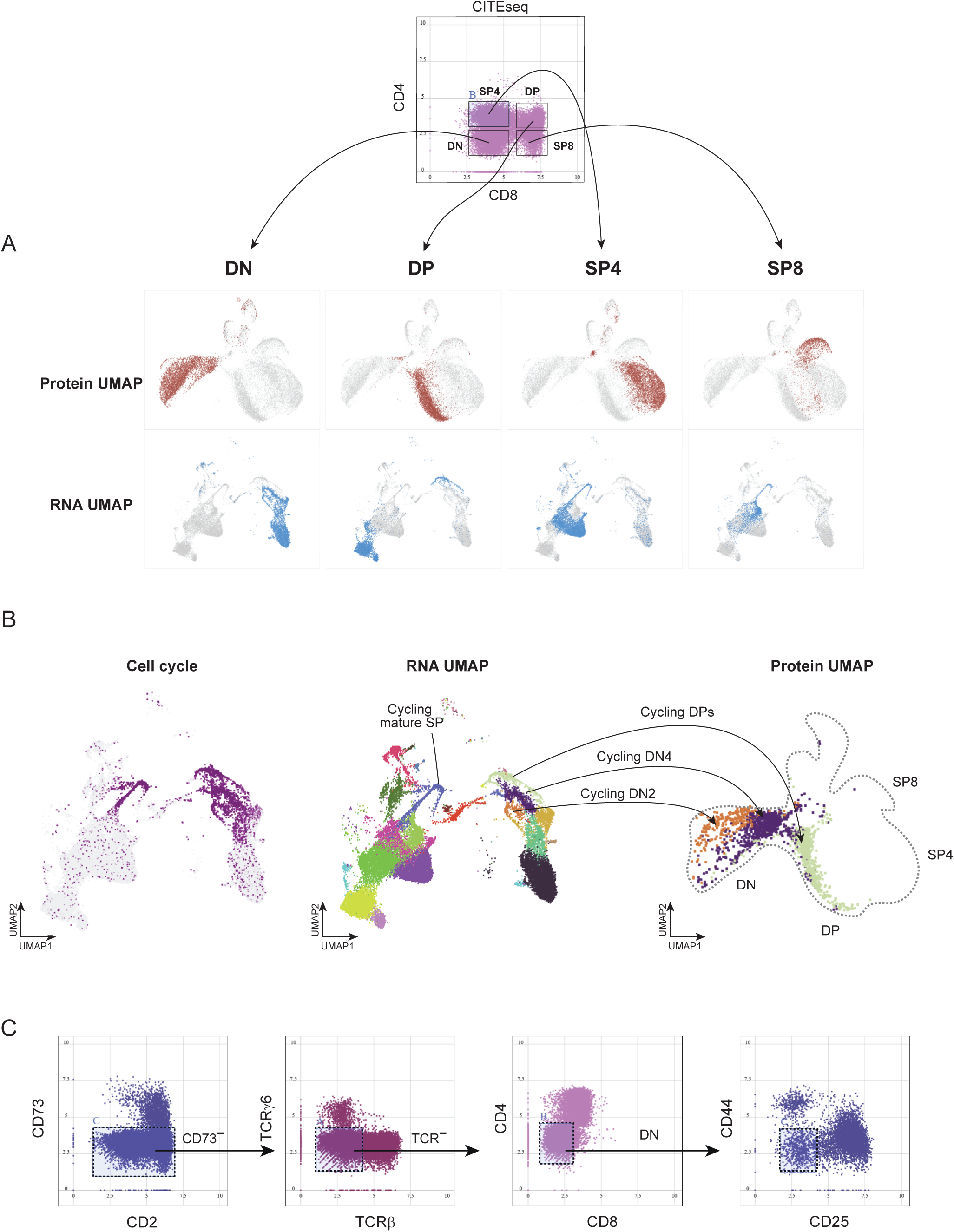
Immature DN thymocytes rearrange TCRα as well as TCRβ. **A.** Definition of thymocyte populations from the RNA and protein profiles in the combined IGT17/18 (total thymocytes, but downsampled to avoid an excess of DPs, and suspect dying-cell clusters filtered out). Top: plot of CD4 vs CD8 CITEseq data used to define main populations. Below: representation of the gated cells in the RNA and protein UMAPs. **B.** Fine definition of the cycling cell clusters: left, expression of cycling-cell genes (representative *Top2a*) in the RNA UMAP (middle panel). The three cycling clusters are projected onto the protein UMAP (right, as in A). The fourth cluster of cyclingcells (cluster 7 in **Fig. 8A**) corresponds to mature SP4 and SP8 cells, which cycle prior to thymic egress. **C.** Sequential cytometry-like gating of cells from the CITEseq data: CD73-negative, TCR-negative, CD4/8-negative DN cells (for **Fig. 8F-K**)

**Fig. S12:**
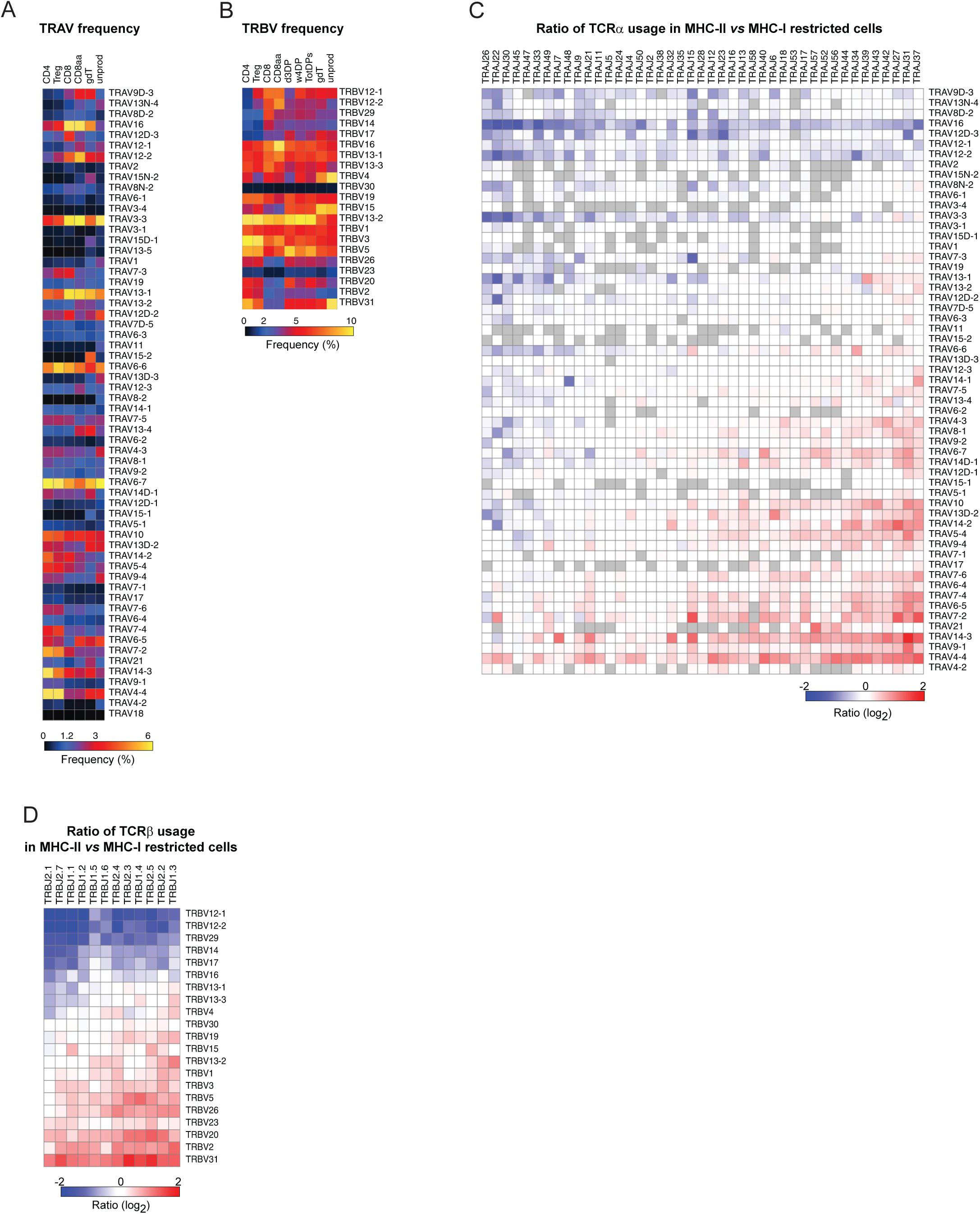
Preference in V gene usage according to MHC restriction. **A.** Heatmap representation of *TRAV* gene usage frequencies (%) in cells of different lineages (CD4, Treg, CD8, CD8aa and gdT), in unproductive rearrangements (all data), ordered by the ratio of averaged frequencies in MHC-I- (CD8 and CD8aa) vs MHC-II-restricted T cells (CD4, Treg). **B.** Similar representation of *TRBV* gene usage frequencies (%) in cells of different lineages: CD4, Treg, CD8, CD8aa, thymic DP (from 3 day-old or 4 week-old mice, or averaged), gdT, and in unproductive rearrangements. **C.** Ratio of usage for all *TRAV/TRAJ* pairs in MHC-II vs MHC-I restricted cells (CD4+ and CD8+, respectively) **D.** Ratio of usage for all *TRBV/TRBJ* pairs in MHC-II vs MHC-I restricted cells (CD4+ and CD8+, respectively).

## EXTENDED DATA TABLE LEGENDS

**Table S1: Characteristics of public clonotypes identified across the immgenT dataset.**

A. List of all public clonotypes identified including their probability of generation (Pgen).

B. List of iNKT and MAIT-related public clonotypes

C. List of Public clonotypes identified predominantly in SFB-positive mice

D. List of Public clonotypes identified predominantly in *Mycobacterium Tuberculosis*-infected mice

E. List of ‘other’ public clonotypes identified

**Table S2:** Characteristics of Treg cells carrying “clumps” of closely related TCRs in the immgenT dataset.

**Table S3:** Characteristics of expanded clonotypes with kidney preferential localization in CD4+ (**A**) and CD8+ (**B**) T cells.

**Table S4:** Details of VDJ rearrangement in the entire non-transgenics immgenT dataset

A. *V* and *J* gene usage frequencies (%) in productive and unproductive *TRA* and *TBB* rearrangements.

B. Number of nucleotides trimmed for each *V* and *J* genes in productive rearrangements.

C. Average number of nucleotides added (N and P) and trimmed in 3’V-GENE and 5’ J-GENE in productive rearrangements.

**Table S5: List and characteristics of cells with unusual rearrangements identified in productive *TRA* and *TRB* rearrangement in non-transgenics ImmgenT dataset.**

A. Cells with two *V* or *J* genes due to the use of cryptic RSS.

B. Cells with inter-gene recombination events.

C. Cells with recurrent, precise off-target events and two J genes in *TRA*.

D. Cells with two *J* genes in *TRA* due to precise off-target RAG activity and hybrid joint.

E. Cells with two *J* genes in *TRB* due to splicing defect.

F. Cells with *TRDD1* and/or *TRDD2* in *TRA*, including rearrangements with *TRDJ1* and *TRAC/TRDC*.

G. Cells with *TRBD1* and *TRBD2* in *TRB* rearrangements.

H. Cells with DNA capture at the junction in *TRA* and *TRB*.

I. Cells with RAG off-target DSB.

J. Cells with trans-rearrangement of *TRAJ* with *TRBD-TRBJ* and of *TRBV* with *TRAJ*.

K. Cells with trans-splicing of the L-PART1 of one gene with the V-EXON of a downstream gene.

L. Number of cells for each trans-splicing event identified.

**Table S6: Characteristics of recurrent rearrangements.**

A. Details of VaJa pairs (AJ) with amino-acids overrepresented junctions in the ImmgenT dataset. VaJa pair of iNKT cell is colored in green and VaJa pair of MAIT cell is colored in orange.

B. Details of VaJa pairs (AJ) with amino-acids overrepresented junctions in human blood T cells.

C. Details of dominant nucleotide sequences in VaJa joins (AJ), including the number of nucleotides added (N) and trimmed (Vdel and Jdel).

**Table S7: Characteristics of recurrent rearrangements in single-chain clonotypes.**

A. Recurring aN-clonotypes in the full dataset.

B. Recurring bN-clonotypes in the full dataset.

C. Recurring bN-clonotypes specific to CD8aa T cells.

D. Recurring aN-clonotypes specific to CD8aa T cells.

E. Recurring bN-clonotypes with the rearrangement *TRBV13-2/TRBJ2-7*.

**Table S8:** Usage frequencies (%) of *V* genes from the productive allele in cells that have an unproductive rearrangement on the second allele, in *TRA* (**A**) and in *TRB* (**B**).

**Table S9:** Characteristics of TCRα clonotypes paired with different TCRβ clonotypes within the same sample.

**Table S10:** *V* gene usage frequencies (%) across different lineages (CD4, Treg, CD8, CD8aa and gdT) and in unproductive rearrangement. The CD4/CD8 ratio was calculated for each *V*-genes in *TRA* (**A**) and *TRB* (**B**). In *TRB* (**B**), *V* gene usage frequencies (%) was also estimated in thymic DP, along with ratios: CD4/Treg, CD4/DP and CD8/DP.

**Table S11:** *V* gene usage frequencies (%) in CD4+ T splenocytes from mice expressing only MHC-II H2-A (A), H2-E (E) or MHC-II deficient (KO), in *TRA* (**A**) and *TRB* (**B**).

**Table S12:** Simplified *TRAV* genes assignment used in repertoire computations.

**Table S13**: SummaryTable of the Human TCR dataset used in the analysis of the recurring rearrangement.

## Supplementary Note

### Artefacts encountered while generating immgenT TCRseq

#### 1. Discordance between read-processing pipelines

The TCRseq data pipeline starts with a sequence assembly step, by computationally aligning and merging overlapping DNA reads to reconstruct the original mRNA. We used the CellRanger VDJ algorithm for this purpose, and all data reported in this paper, in the SummaryTable and the GEO submissions stems from CellRanger processing. In some circumstances, however, when we wanted to verify that the contig generation algorithm was not generating artefactual structures, we compared outputs from CellRanger and from an independent processing suite, MiXCR (v4.7.0) ^52^. For the most part, there was good agreement with the output from the two pipelines (>90% concordance), especially when dealing with high-quality datasets. On the other hand, even after stringent thresholds on read quality (e.g. CellRanger cell calling criteria, RNA/ADT count filtering and singlets filtering), we still observed some degree of discordance between the two outputs, as illustrated in Fig. A for *TCRa* and *TCRb* from IGT19,. Some discordance corresponded to contigs assembled by one pipeline but not the other (6 % exclusive to CellRanger, 5% exclusive to MiXCR for *TRA*, see Fig. A; 3% exclusive to Cellranger, 1% exclusive to MiXCR for TRB, see Fig. A).

Perhaps even more disquieting, however, was that in those cells in which a contig was called by both the CellRanger and MiXCR pipelines, the actual TCRs found were not the same in ∼4% of cases (both sets of contigs evaluated by IMGT-VQUEST with the same B6 reference library) even after switching between beta1 and beta2 (alpha1 and alpha2) the assemblies assigned as primary or secondary. The different calls were usually provoked by one differing element (e.g. TRBV and TRBJ matched but the junction was different, or junction and TRBJ matched but with a different TRBV assigned).

#### 2. First and second outputs from CellRanger processing are the same for some cells

In 747 cases over the entire dataset, the CellRanger pipeline generated two quasi-identical contigs as beta and beta2 (and 407 for alpha and alpha2), which V-QUEST then assigned to the same vgene, j.gene and junction.nt. On examination, these called contigs differed at their extremities, but were identical in the V, junction and J regions. The duplicative alpha2 and beta2 information was removed from the final TCR SummaryTable and before analyses reported here.

#### 3. Barcode multiplex artefact

This artefact of the single-cell microfluidic process has been well documented in the case of single-cell ATACseq ^85^, but to our knowledge has not been considered in microfluidics-based TCRseq studies. Although the relative abundance of cells and beads during encapsulation in the 10X Genomics Chromium is tailored such that cells are mostly lysed in a drop together with only one bead, it does happen that two beads (each of which carries a unique cell barcode) are present together with one cell. After library construction and data processing, the result appears as two cells with the exact same TCR and highly related gene expression profiles.

**Fig. A.**
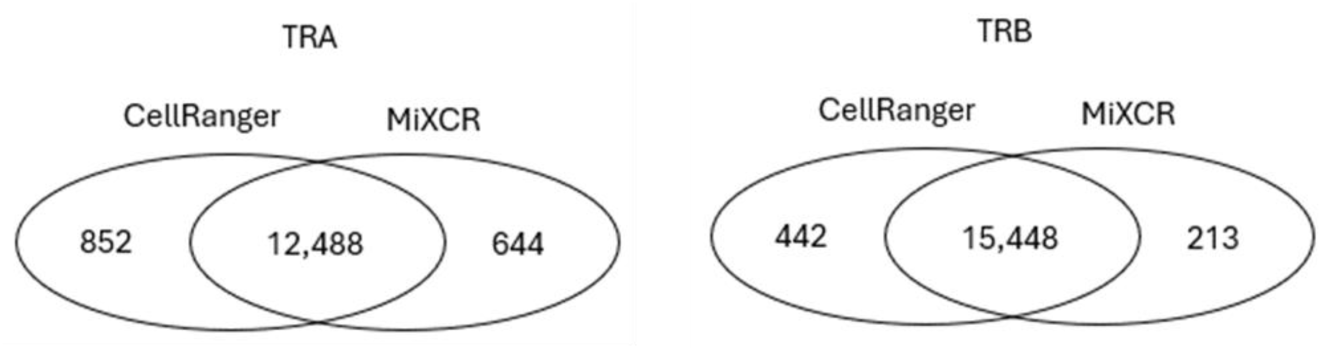
Comparison of contig assembly between CellRanger and MiXCR (IGT19). Venn diagrams reporting the number of cells yielding a contig for TCRa or TCRb (left, right) with CellRanger or MiXCR processing (same B6 reference used for both). While both algorithms similarly identified contigs in the majority of cells, 10.7% and 4.15 of cells only yielded a contig with only one of them for TCRa and TCRb, respectively.

To evaluate this issue, we analyzed results from experiments in which the same sample was run on two independent lanes (Fig. B). In the majority of cases, inter-lane duplicates were roughly as frequent as intra-lane duplicates, implying that there was no generalized problem. Note that the variation in duplication frequency for the different IGT pairs is expected, low in the thymus (IGT17/18) much larger in low-complexity intestinal tissue (IGT20/21). The one standout was in thymic double-positive cells of IGT72, with 57 duplicates that were not matched in the corresponding IGT71. In this case, a technical problem may have occurred which led to more frequent barcode multiplex in the IGT72 run.

In practice, because the problem seemed infrequent and of very low proportion (affecting only 0.004 of cells in IGT72) we did not consider it further, except when the identity of a specific TCR clonotype was important: in those cases, we consider duplicates for followup action only if confirmed by the other lane.

#### 4. Suspect inter-dataset duplications

Some clonotypes (public specificities) are expected to be repeated in different datasets, sometimes even with identical junction sequences at the nucleotide level. But we also observed a sizeable number (1223) of such duplications that also corresponded to identical cell barcodes (which should be a vanishingly rare occurrence, given the >10^6^ cell barcodes in each 10X run). Such events most likely stem from contamination during the library preparation, or “index hop-ping” during sequencing (plausible because such events occurred only between datasets whose TCR libraries were processed and sequenced together). These were systematically eliminated from the SummaryTable and the analyses reported here.

**Fig. B.**
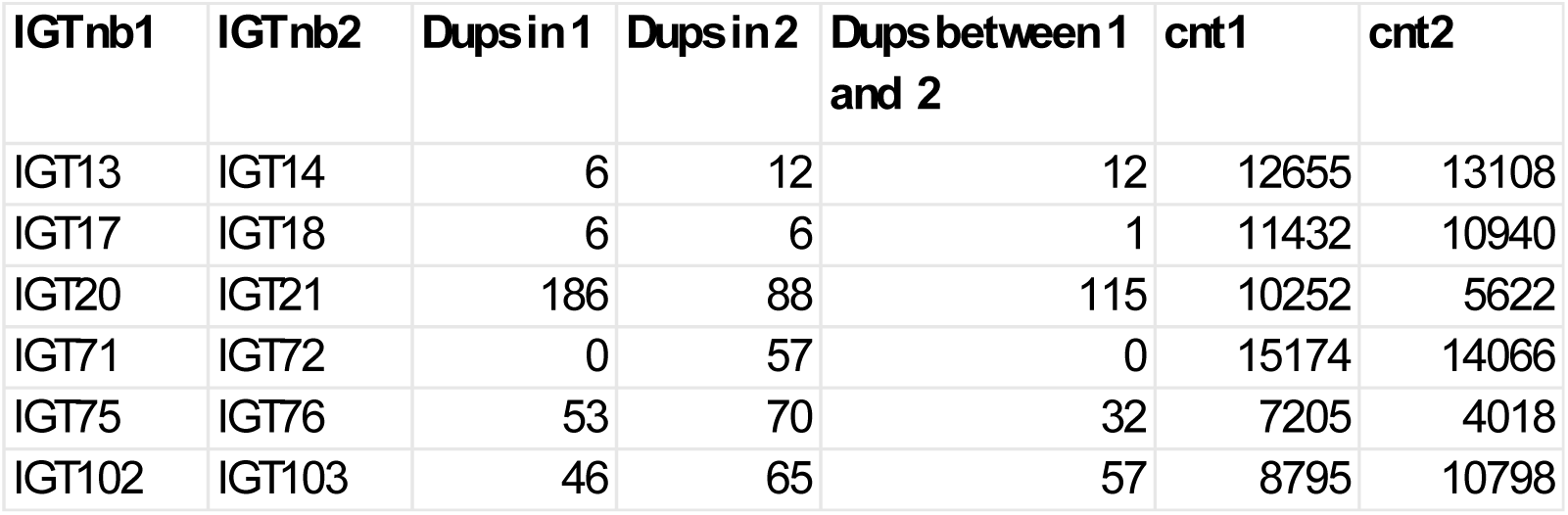
Evaluation of inter vs intra-batch duplicates to find possible “barcode multiplex” artefacts. These artefacts are difficult to evaluate in single-cell TCRseq data, since duplication from clonal expansion is expected, especially in some low-complexity samples. However, it is possible to evaluate its incidence when one cell sample is split into two independent lanes of the Chromium instrument: if duplicates are true, they should appear as frequently between the two lanes as within a single lane. Barcode multiplex events, on the other hand, yield duplicates only within a single lane. For all IGT datasets for which the same cell sample was split into two independent lanes of the Chromium instrument, and encompassed sizeable numbers, the figure shows the number of duplicated (n=2) clonotypes within one or the other IGT lane, and between the two lanes. Total cell numbers in the two lanes are shown at right.

#### 5. Ambient RNA contamination

Single-cell technologies based on microfluidic encapsulation of cells with barcoded beads are prone to the “ambient RNA” artefact: during the last stages of cell preparation, damaged or dying cells release RNA into the medium. Subsequently, the microdrop that carries a cell also takes in a few random molecules of this soluble RNA, which then end up incorporated into the cell’s mRNA or TCR libraries. In principle, this contamination could particularly become an issue for TCRseq, pick up of stray TCRa or TCRb transcripts giving the appearance of a cell with several TCR transcripts. To some extent, this is dealt with by the CellRanger pipeline, which drops contigs generated from few reads, but is not null.

To evaluate this issue in immgenT TCR data, we followed two approaches

**Table A:** ambient TCR RNAfrequency in empty beads.

| Dataset | Nb of Empty Beads | Nb of beads with contig | Nb of beads with VaJa | Nb of beads with VbJb | Nb beads with both | Overall Va % | Overall Vb % |
| --- | --- | --- | --- | --- | --- | --- | --- |
| IGT7 | 694119 | 6044 | 1637 | 3733 | 113 | 0.24 | 0.54 |
| IGT10 | 720526 | 2163 | 519 | 1368 | 12 | 0.07 | 0.19 |
| IGT12 | 720752 | 3440 | 720 | 2089 | 24 | 0.1 | 0.29 |
| IGT17 | 712306 | 10377 | 1151 | 5985 | 122 | 0.16 | 0.84 |
| IGT33 | 701491 | 1450 | 410 | 787 | 5 | 0.06 | 0.11 |
| IGT36 | 712271 | 215 | 12 | 46 | 0 | <0.01 | 0.01 |
| IGT67 | 612118 | 1530 | 299 | 975 | 4 | 0.05 | 0.16 |
| IGT74 | 711506 | 1009 | 262 | 577 | 4 | 0.04 | 0.08 |
| IGT85 | 622433 | 6393 | 1561 | 3924 | 111 | 0.25 | 0.63 |
| IGT88 | 635653 | 6847 | 1196 | 4106 | 219 | 0.19 | 0.65 |

a. Empty beads. In droplet-based single-cell RNA-seq, droplets containing a barcode-carrying bead but no actual cell (“empty beads”) do contain ambient RNA. Empty beads are readily identified in the data processing (cell barcodes with very low UMI and gene counts) and are normally eliminated. Here, we recovered and analyzed the TCR sequences generated from empty beads. As shown in Table A, the frequency of ambient RNA detection varied with the datasets, likely as a function of the difficulty of cell preparation and hashtagging. TCRb was found more than TCRa, reflecting the different transcript levels in most T cells. Predictable, empty beads with both TCRa and TCRb were quite rare. The final columns on TableA show that the incidence of TCRa or TCRb contamination can range as 0.01 to 0.25% and 0.01 to 0.84%, respectively, which should be taken into consideration when dealing with rare events.
b. Datasets with TCR transgenic samples. Some IGT datasets contained cells from TCR transgenic mice, as well as sapes from non-transgenic animals (in particular the constant Control Spleen samples), all from normal B6 mice. Ambien RNA would thus appear as discovery of transgenic clonotypes in non-transgenic samples. In combined IGT95/96, we detected 7 instances of the OT-II TCRb chain among 485 control cells (1.44%). In IGT40, we detected 3 instances of the Smarta or P14 TCRa or TCRb chain among 1297 cells (0.23%), thus in range with the estimates from empty beads.

#### 5. Apparent mutations

Somatic hypermutation (SHM) occurs in B cells at high rates in rearranged Immunoglobulin genes during affinity maturation in germinal centers^86^, preferentially targeting the CDRs. SHM is generally thought not to occur in T cells, except in some species like nurse sharks^87^. Because immgenT datasets covered many exotic states and locations, particularly the germinal center, it seemed a valuable context to search for SHM, in case it occurred in peculiar states or locales for mouse T cells. We used very stringent processing criteria to avoid artefacts from PCR or sequencing errors (only high-quality reads were retained, only productive full-length contigs with >1 Unique Molecular Identifiers (UMI) were considered). We observed a low mutation rate (2×10^-4^ to 2×10^-5^ mutations per 100 bp), not very different between V and C regions, and with no preferential relationship to organ or T cell state. We did observe a number of intriguing recurring at the same locations, and enriched in *TRAV*-FR3 (Fig. C Table S14). We also observed one mutation at position 40 of *TRBJ1-5*, present in a large proportion of cells (553 of 5,893) in 24 different samples of B6 mice of JAX origin, with a variable proportion of mutant-vs WT-expressing cells. However, these recurring mutations were often detected in UMIs that also yielded reads with the WT sequence, strongly suggesting PCR artifacts recurring at the same position, presumably because secondary structure in the RNA leads to preferential misincorporation by the Taq polymerase used in the PCR steps. Moreover, neither the TRAV-FR3-enriched mutations nor the *TRBJ1-5* mutation at position 40 were detected in the Parse spleen dataset. This discrepancy, combined with the presence of mixed WT/mutant reads in the raw data, further supports the interpretation that these mutations are PCR artefact rather than SHM.

**Fig. C.**
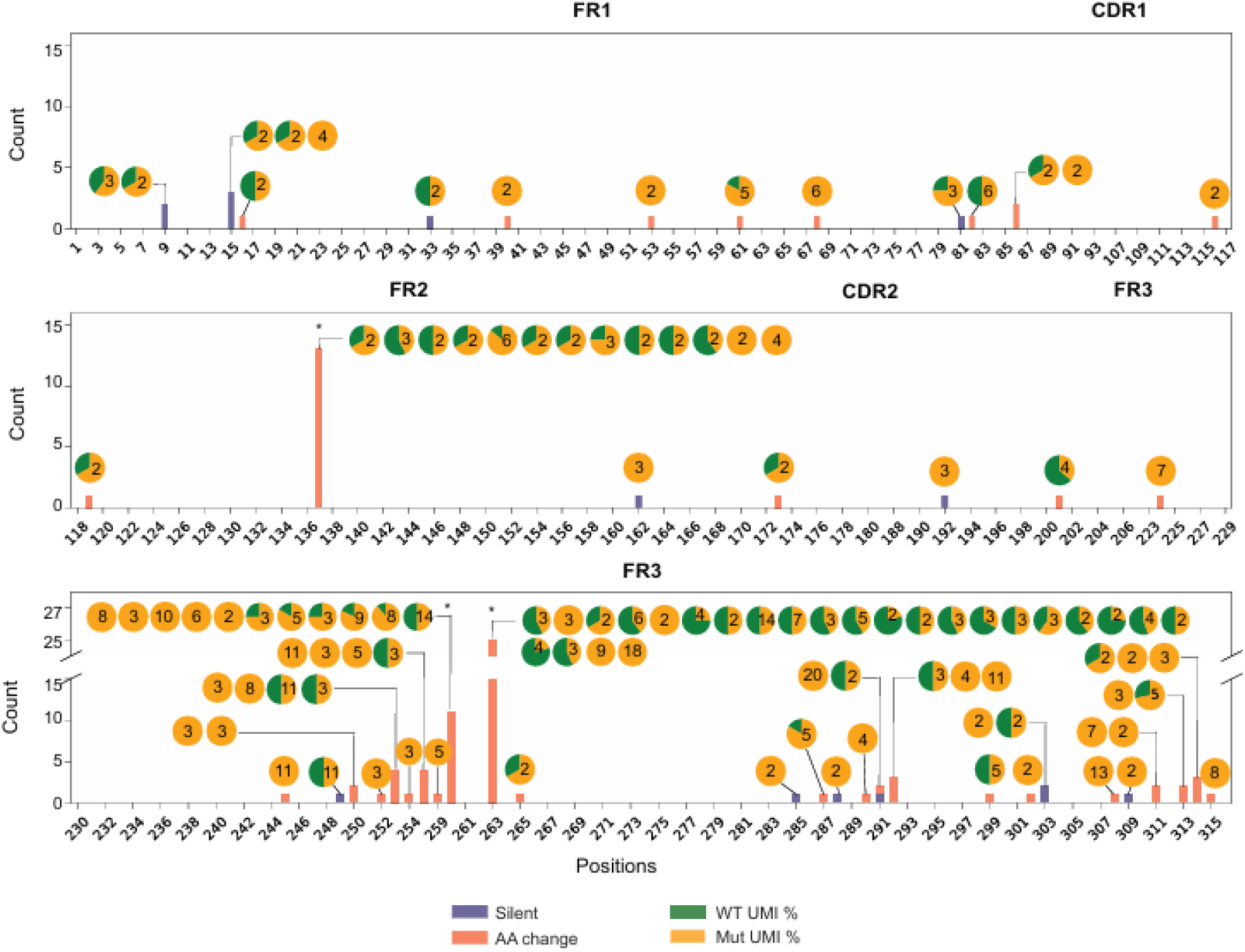
Recurring mutations in TRAV, most likely artefactual.

